# Rapaprotin is Activated by an Endopeptidase to Disassemble 26S Proteasome

**DOI:** 10.1101/2025.02.21.639179

**Authors:** Hanjing Peng, Zufeng Guo, Wei Li Wang, Deyao Yin, Shitao Zou, Thomas Asbell, Brett R. Ullman, Maya Thakar, Feiran Zhang, Sam Y. Hong, A. V. Subba Rao, Kunyu Wang, Shuwen Zhang, Zhaolong Wu, Xuemei Li, Seth S. Margolis, William H. Matsui, Christian B. Gocke, Youdong Mao, Jun O. Liu

## Abstract

The 19S regulatory particle (RP) associates with the 20S core particle (CP) to form the 26S proteasome, an evolutionarily conserved holoenzyme that plays key roles in both physiological and pathological processes. Proteasome inhibitors that target the catalytic subunits within the 20S have proven to be valuable research tools and therapeutics for various cancers. Herein we report the discovery of rapaprotin, a 26S proteasome assembly inhibitor from our natural product-inspired hybrid macrocycle rapafucin library. Rapaprotin induces apoptosis in both myeloma and leukemia cell lines. Genome-wide CRISPR-Cas9 screen identified a cytosolic enzyme, prolyl endopeptidase (PREP) that is required for the pro-apoptotic activity of rapaprotin. Further mechanistic studies revealed that rapaprotin acts as a molecular transformer, changing from an inactive cyclic form into an active linear form, rapaprotin-L, upon PREP cleavage, to block 26S proteasome activity. Time-resolved cryogenic electron microscopy (cryo-EM) revealed that rapaprotin-L induces dissociation of the 19S RP from the 26S holoenzyme, which was verified in cells. Furthermore, rapaprotin exhibits a marked synergistic effect with FDA-approved proteasome inhibitors and resensitizes drug-resistant multiple myeloma cells from patients to bortezomib. Taken together, these results suggest that rapaprotin is a new chemical tool to probe the dynamics of the 26S proteasome assembly and a promising anticancer drug lead.

## Introduction

The 20S core particle (CP) of the many cellular proteasome complexes is an evolutionarily conserved machinery that plays an essential role in protein degradation in eukaryotes, archaea and a subset of bacteria (*1, 2*). When associated with the 19S regulatory particle (RP), the 20S CP transforms into a 26S complex that is essential for all ubiquitin-mediated proteasome-dependent protein breakdown. The function of the 26S proteasome is vital for regulating a myriad of cellular processes, including signal transduction, cell cycle, antigen presentation, apoptosis, and tumorigenesis (*2-5*). Through decades of research, highly selective inhibitors of the three catalytic β-subunits within the 20S CP have been developed and are currently used in the clinic to treat patients suffering from various forms of cancer. The rationale to target proteasome for developing anticancer drugs was initially predicated on the notion that the 26S proteasome is required for the activation of the anti-apoptotic transcription factor NF-κB through degradation of ubiquitinated IκB (*6*). It was discovered that proteasome inhibition is exceptionally effective at killing plasma cells, which are reliant on the proteasome to dispose of misfolded immunoglobulins (*7*). Currently, three proteasome inhibitors, bortezomib (*7, 8*), carfilzomib (*9*) and ixazomib (*10, 11*), have been approved for clinical use and remain primary prescriptions for treatments of multiple myeloma (MM) and mantle cell lymphoma (*12*). Aside from MM and lymphoma, proteasome inhibitors have also shown promise in treating a number of chronic disorders including autoimmune diseases and organ rejection (*13-16*).

Despite the success of the existing proteasome inhibitors, patients undergoing targeted proteasome therapy eventually succumb to drug resistance (*10, 17*). Moreover, these proteasome inhibitor therapies have major side effects (*10*), including peripheral neuropathy, fatigue, thrombocytopenia, and anemia. Although carfilzomib exhibits reduced peripheral neuropathy, it retains the majority of the other side effects of bortezomib and ixazomib, and has additional cardiotoxicity (*12, 18*). These shared side effects among different proteasome inhibitors seem to be mechanism-based and cannot be reduced through alteration of their subunit selectivity. Given the significant clinical benefit of targeted proteasome therapies and the emerging limitations of this approach, agents with novel mechanisms of action that overcome proteasome inhibitor drug resistance with reduced side effects are of critical importance. Herein we report the discovery of a first-in-class proteasome assembly inhibitor named rapaprotin that, in contrast to known proteasome inhibitors, acts by inducing 26S proteasome disassembly, exhibits greater selectivity towards MM cells over normal endothelial cells, and is synergistic with existing proteasome inhibitors, being capable of resensitizing patient-derived, drug-resistant MM cells to bortezomib.

## Results and Discussion

### Identification of rapaprotin as a selective inhibitor of MM

We have previously developed a new class of hybrid macrocycles called rapafucins that were inspired by the unique scaffolds and extraordinary mode of action of the natural products-derived drugs rapamycin and FK506 (*19, 20*). We borrowed the FKBP-binding domain of rapamycin and replaced its mTOR-interacting domain with a new combinatorial oligopeptide library. To identify inhibitors of MM, we screened the rapafucin library using the NCI-H929 cell line in a resazurin viability assay (Figure 1a, Figure S1a). The rapafucin library was made by a split-and-pool synthesis and contained a total of 45,000 compounds in 3000 individual pools, with each pool consisting of 15 rapafucins that differ in the first amino acid building block in the effector domain (*19, 20*). The library was screened at a fina concentration of 200 nM per compound or 3 µM total rapafucin. The initial screen yielded 40 hits (Figure 1a, Figure S1a, B); re-evaluation and prioritization based on their dose-dependent activities in a variety of cancer cell lines (NCI-H929, Jurkat-T, A549 and HCC1954) narrowed them down to 4 MM-selective hits (Figure S1b). The top four pools (E26, E27, E28, and A858) that showed 99.7% inhibition in MM viability assay were selected for further deconvolution and validation. Thus, each of the 15 individual compounds from the pools was synthesized and tested in the cell viability assay. Rescreening of the decoded individual rapafucins from Pool E27 (Figure S1c) gave rise to the most potent hit E27-9 with an IC_50_ of 468 nM (Figure 1a, b).

**Figure 1.**
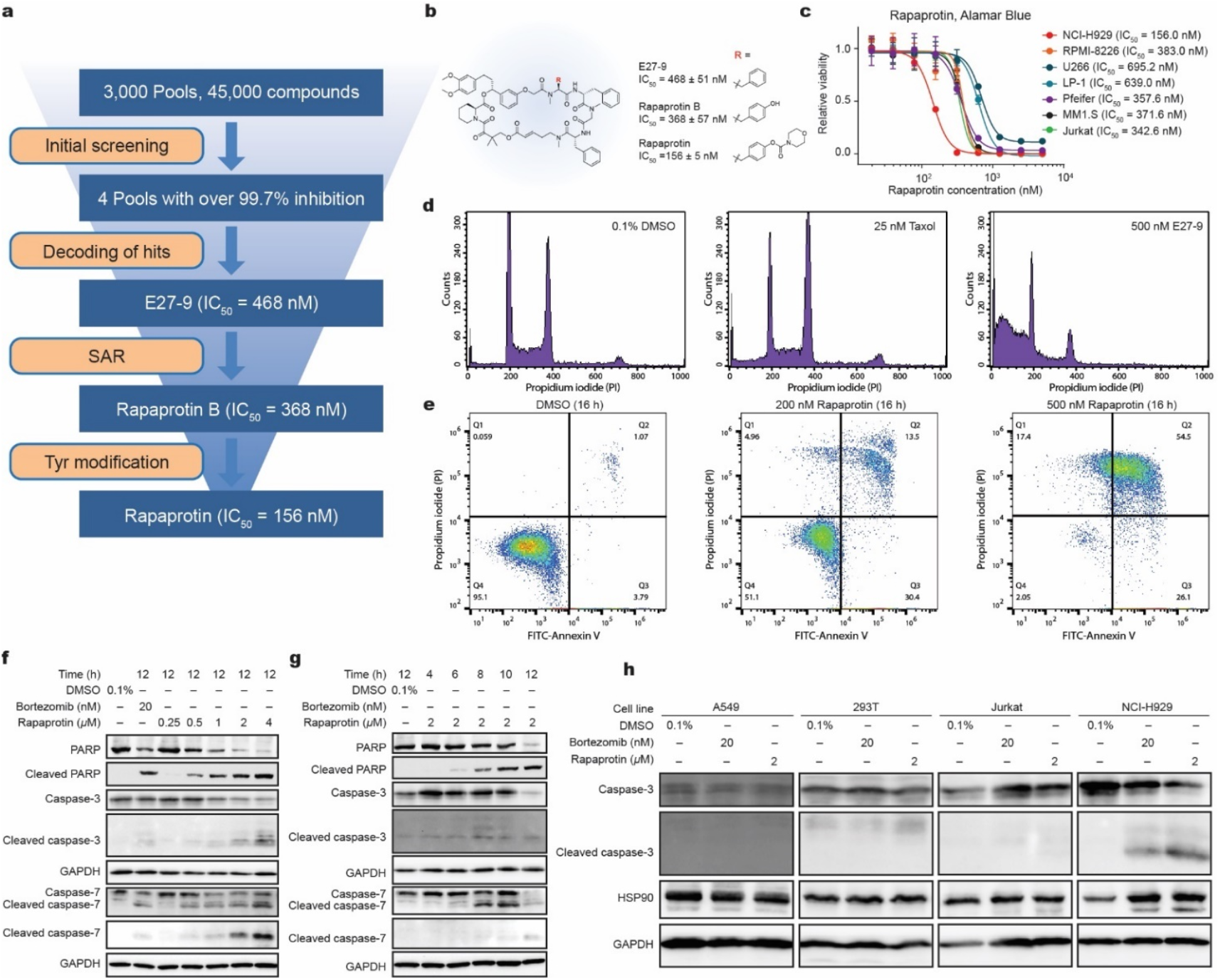
Identification of rapaprotin as selective apoptosis inducer in multiple myeloma cells. (a) Flow chart of anti-myeloma screening, hits decoding and optimization. (b) The structure of rapaprotin, and its analogues E27-9 and rapaprotin B. (c) Dose-dependent cytotoxicity of rapaprotin in select myeloma and lymphoma cells as measured with resazurin viability assay. (d) Cell cycle analysis shows sub-G1 population after E27-9 treatment for 16 hours at 500 nM as measured by flow cytometry with propidium iodide (PI) after fixing the cells. (e) Rapaprotin dose-dependently induces apoptosis in NCI-H929 as measured by flow cytometry with FITC annexin V and PI. (f) Western blot shows dose-dependent activation of caspases 3 and 7 by rapaprotin in NCI-H929. (g) Western blot shows time-dependent activation of caspases 3 and 7 in NCI-H929 cells. (h) Western blot shows 8-hour rapaprotin treatment leads to activation of caspase-3 in NCI-H929, but not in A549, 293T, or Jurkat-T cells. Please refer to Supporting Information for uncropped blots for Figure 1f, g and h.

To further improve the potency of E27-9, we conducted a structure-activity relationship (SAR) study by systematically altering each of the four amino acid building blocks in the effector domain of E27-9. Among a number of analogs synthesized and tested, an analog with a slightly improved potency (IC_50_ = 368 nM), named rapaprotin B, emerged upon replacement of a phenylalanine in E27-9 with a tyrosine residue (Figure 1b). Modification of the tyrosine hydroxyl group with a morpholine-containing moiety yielded rapaprotin with greater solubility than rapaprotin B (Figure S1d) and a significantly improved potency (IC_50_ = 156 nM) (Figure 1b). Aside from NCI-H929, we also assessed the effects of rapaprotin on several other MM and hematopoietic cancer cell lines. Though NCI-H929 is most sensitive to rapaprotin, other MM cell lines are also inhibited by rapaprotin, albeit with higher IC_50_ values (Figure 1c).

### Rapaprotin induces apoptosis through activation of caspases in MM cells

To further delineate the cellular effects of rapaprotin on MM cells, we determined the changes in cell cycle distribution of NCI-H929 cells upon treatment with rapaprotin. After exposure to E27-9 (the initial rapaprotin lead) for 16 h, there was a significant accumulation of sub-G_0_ cell population at the expense of cells in the G1, S and M phases, suggesting the induction of apoptosis (Figure 1d). As a positive control, the microtubule inhibitor taxol caused cell cycle arrest in the G2/M phase as expected. The rapaprotin-induced apoptosis of NCI-H929 cells was confirmed by surface expression of annexin V and propidium iodide staining (Figure 1e). Trypan Blue cell counting experiment also shows massive cell death after 14 hours of treatment with rapaprotin (Figure S1e).

We next determined the effect of rapaprotin on the activation of caspases. Indeed, rapaprotin induced the cleavage of procaspases-3, -7 and -8 in a dose-dependent manner (Figure 1f, S1f). Procaspase cleavage was also time-dependent, with procaspase-3 and -7 cleavage occurring 8-10 h after treatment with 2 µM rapaprotin (Figure 1g). Rapaprotin, like bortezomib, caused significant cleavage of pro-caspase-3 in NCI-H929 and to a lesser degree in Jurkat T cells. In contrast, it failed to cause pro-caspase cleavage in HEK293T kidney epithelial cells or A549 lung cancer cells (Figure 1h), consistent with the lack of inhibition of those cell lines by rapaprotin (Figure 1c, Figure S1g). Consistent with the caspase activation, rapaprotin also induced PARP cleavage in a dose- and time-dependent manner in NCI-H929 cells (Figure 1f, g).

### Activation of rapaprotin by prolyl endopeptidase

As rapaprotin induces apoptosis in MM cells, we speculated that a loss-of-function screen to select rapaprotin-resistant mutant clones would reveal essential mediators of its cellular activity. We thus conducted a genome-wide CRISPR-Cas9 knockout screen using the Brunello library with sgRNAs that target 19,114 different human genes (Figure 2a). Pooled NCI-H929 cells transduced with the library were cultured with rapaprotin at such concentrations that, on average, approximately 1-5 in every one-million cells survived and formed colonies (Figure S2a). A total of 55 surviving colonies were isolated, expanded and reassessed for resistance to rapaprotin. Fourteen of the 55 colonies exhibited persistent resistance to rapaprotin by the viability assay while remaining sensitive to bortezomib (Figure 2b and Table S1). Sequencing analysis of the rapaprotin-resistant single-cell clones revealed that 11 out of the 14 cell lines harbored sgRNAs targeting the gene encoding prolyl endopeptidase (PREP) (Table S1). In contrast to the parent NCI-H929 cells, a clone harboring the PREP sgRNA was refractory to rapaprotin-induced PARP cleavage and caspase-3 activation while remaining sensitive to bortezomib (Figure 2c, d). To confirm that PREP was indeed required for the proapoptotic activity of rapaprotin, we generated multiple additional CRISPR knockout lines of NCI-H929 using four different sgRNAs against PREP. In comparison to parental NCI-H929 cell line or the control line using a scrambled non-targeting (NT) sgRNA sequence, all four PREP knockout sublines were completely resistant to rapaprotin (Figure 2e, f). Furthermore, a highly potent and specific inhibitor of PREP, S-17092, also conferred resistance to rapaprotin in a dose-dependent manner (*21*) (Figure 2g). Together, these results confirmed that PREP is required for the proapoptotic activity of rapaprotin, suggesting that rapaprotin may be activated by PREP in MM cells.

**Figure 2.**
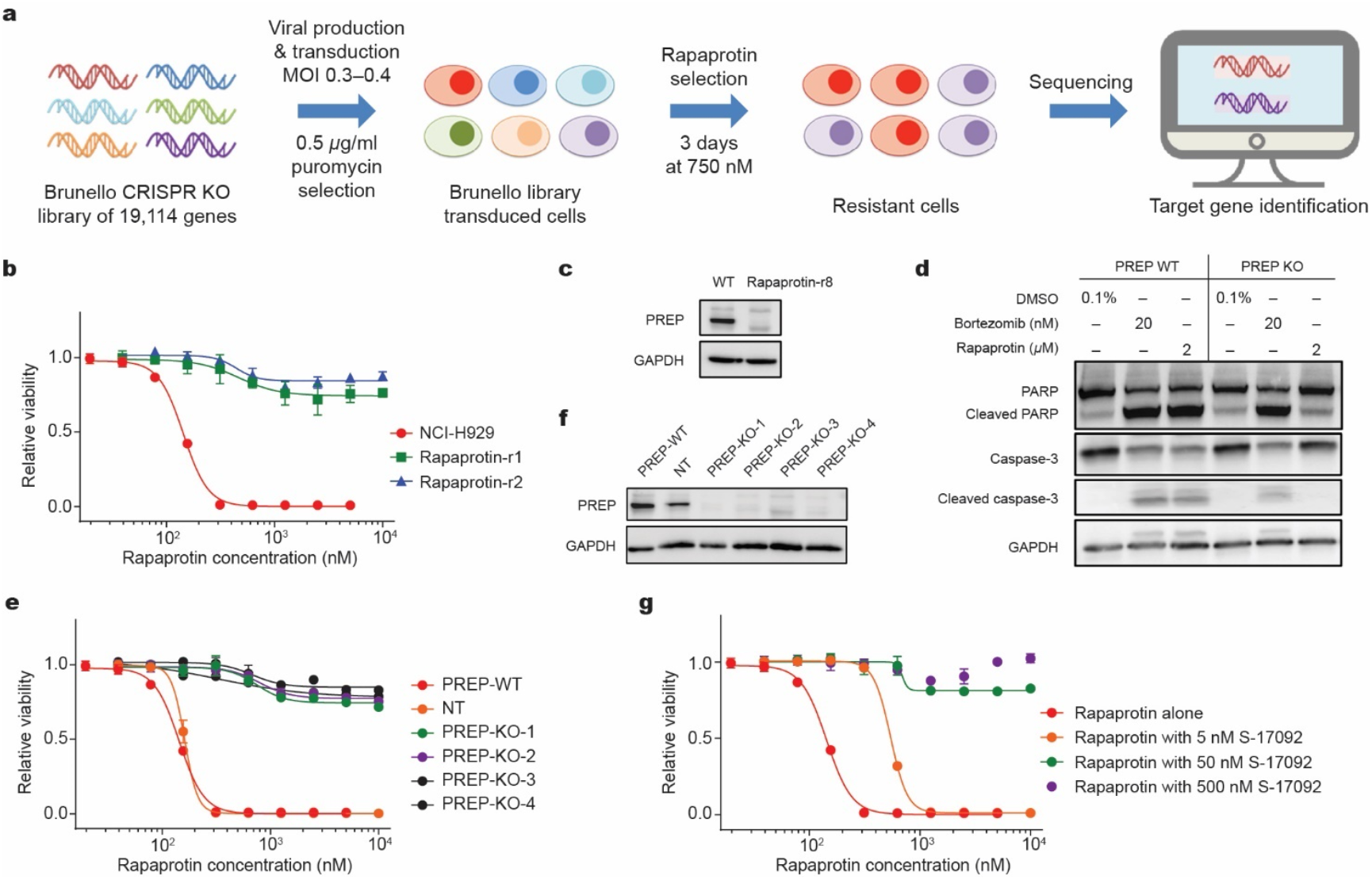
Rapaprotin is activated by PREP-catalyzed cleavage. (a) Flow chart of CRISPR screening to identify PREP as a key factor for rapaprotin activity. (b) Confirmation of rapaprotin resistance of cell lines resulted from CRISPR screening, Rapaprotin-r1 and Rapaprotin-r2 as examples, as determined by 72-hour resazurin viability assay. (c) Confirmation of complete knock-out of PREP, Rapaprotin-r8 as an example, as determined by Western blot. (d) Western blot shows activation of PARP and caspase 3 in WT NCI-H929 but not in PREP KO cells. (e) Knocking out PREP in WT NCI-H929 results in complete resistant to rapaprotin. (f) Western blot shows complete knock-out of PREP in NCI-H929 cells. (g) PREP inhibitor S-17092 reverses cytotoxicity of rapaprotin in NCI-H929, as determined by 72-hour resazurin viability assay. Please refer to Supporting Information for uncropped blots for Figure 2c, d and f.

PREP belongs to the serine peptidase family and is known to cleave proteins and peptide substrates (*22*). Rapaprotin contains multiple amide and ester bonds, rendering it a potential substrate of PREP. To determine whether rapaprotin is hydrolyzed by PREP, we incubated rapaprotin with recombinant PREP enzyme in vitro and followed the reaction using liquid chromatography-mass spectrometry (LC-MS). Upon incubation with PREP overnight, the cyclic rapaprotin is converted into a new product (Figure 3a, b). In contrast, there was only a small amount of hydrolysis of cyclic rapaprotin in the same buffer in the absence of PREP. The hydrolytic product generated by PREP has a shorter retention time in LC chromatogram (Figure 3b), suggesting that it has higher polarity. It has a molecular mass of 1351.6 that is 18 Dalton heavier than the cyclic rapaprotin (MW: 1333.6), which is indicative of addition of a water molecule. We thus named it rapaprotin-L because hydrolytic cleavage of cyclic rapaprotin would give rise to a linear product.

**Figure 3.**
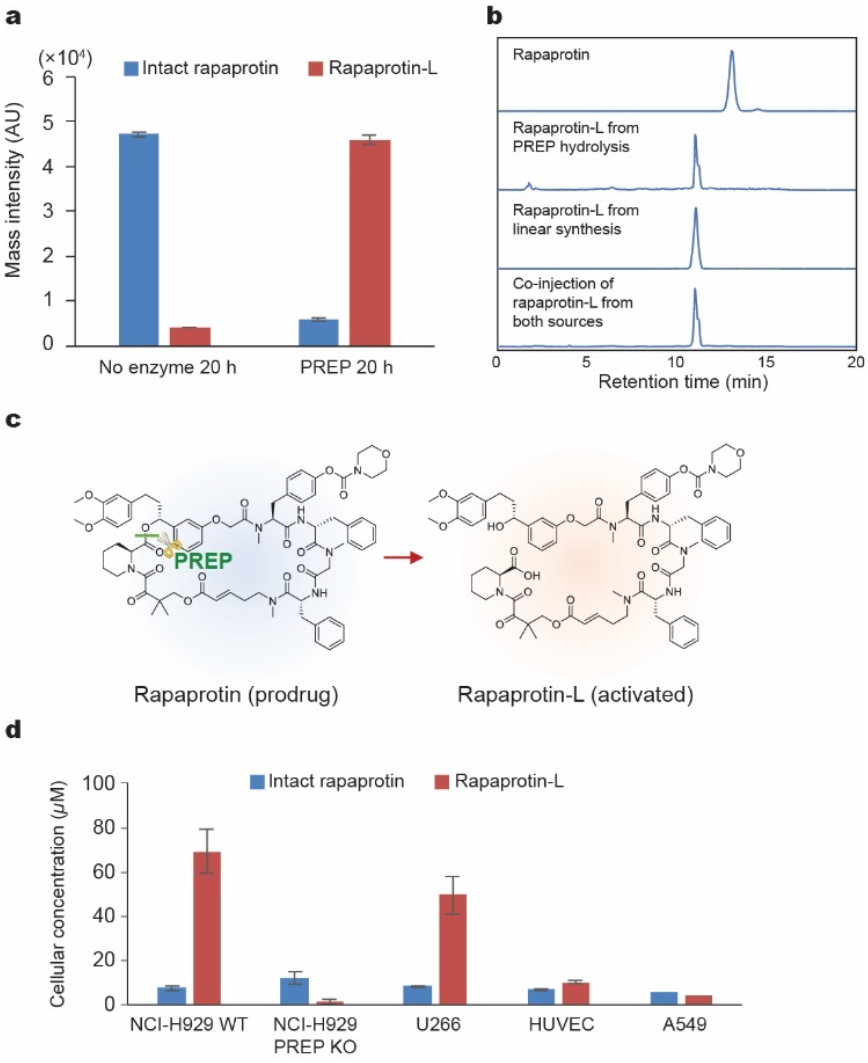
Rapaprotin-L is generated through PREP mediated cleavage. (a) Rapaprotin is hydrolyzed to rapaprotin-L by recombinant PREP. (b) HPLC-MS of rapaprotin and rapaprotin-L from PREP hydrolysis, linear synthesis, and co-injection of rapaprotin-L from both sources. (c) Structure of rapaprotin and rapaprotin-L. (d) Cellular concentration of rapaprotin and rapaprotin-L in different cell lines.

To confirm that PREP cleaves rapaprotin in cells, we incubated NCI-H929 and rapaprotin-resistant PREP-KO cells, respectively, with 1 µM of rapaprotin for 3 hours and detected cyclic rapaprotin and rapaprotin-L in cell lysates. Rapaprotin-L was observed in WT NCI-H929 but not in PREP-KO cells (Figure S2B). We next used LC-MS to determine the concentration changes of intracellular rapaprotin-L over a time course of 24 hours and found dramatic increase of rapaprotin-L concentrations (over 120 µM after 8 hours of incubation) in wild-type but not PREP-KO cells (Figure S2c).

PREP is known to cleave polypeptides with proline at the P1 position (*23*). Although rapaprotin does not contain a proline residue, the pipecolate moiety within the FKBD domain is a close mimic of proline residue. We thus hypothesized that the pipecolate-containing lactone might be the site of cleavage by PREP (Figure 3c). To test this hypothesis, we undertook a solid-phase total synthesis of the proposed rapaprotin-L (Figure S3). Starting with the lower fragment of FKBD **1** tethered to solid beads to form intermediate **2**, we coupled each of the four amino acid building blocks of the effector domain to give rise to the tetrapeptide-containing intermediate **6**. Upon deprotection of Fmoc protecting group, **6** was coupled to the upper fragment of FKBD **7** to yield intermediate **8** on beads. Detachment of **8** from solid phase followed by Noyori asymmetric reduction of the released intermediate **9** gave the final product rapaprotin-L, which was confirmed by HPLC and high-resolution MS.

We next compared the LC-MS profile of the synthetic rapaprotin-L with that of the hydrolytic product generated from incubation of cyclic rapaprotin with recombinant PREP enzyme (Figure 3b). The retention time of the PREP-generated hydrolytic product is nearly identical with that of synthetic rapaprotin-L as demonstrated by co-injection of the two samples (Figure 3b). Together, these results strongly suggest that rapaprotin undergoes PREP-catalyzed hydrolysis to generate rapaprotin-L. Given that the PREP enzyme is required for the proapoptotic activity of rapaprotin, it is likely that rapaprotin-L, rather than cyclic rapaprotin, is the active species inducing apoptosis. In support of this notion, we observed accumulation of rapaprotin-L in WT NCI-

H929 and U266 cell lines but not in PREP-deficient, rapaprotin-resistant clones or rapaprotin-insensitive HUVEC or A549 cells most sensitive cell line (IC_50_ = 143 nM), has the highest intracellular rapaprotin-L concentration (69 µM). U266, which is less sensitive to rapaprotin (IC_50_ = 695 nM), also has lower rapaprotin-L concentration (49 µM). As expected, significantly lower intracellular rapaprotin-L was detected in HEK293T (18.8 µM), A549 (4.2 µM) and E2R12 (1.4 µM, PREP KO cells), which are insensitive to rapaprotin, further supporting that rapaprotin-L is the active species causing cell death.

### Rapaprotin inhibits the ubiquitin-proteasome pathway

The ability of rapaprotin to induce apoptosis and activate caspases in MM cell lines is reminiscent of proteasome inhibitors (*24*). We thus determined whether rapaprotin affected the ubiquitin-proteasome pathway using a destabilized dihydrofolate reductase mutant (mDHFR)-GFP reporter (*25*) (Figure 4a). Upon ectopic expression, the mDHFR-GFP fusion protein undergoes constant ubiquitination and consequent degradation by the proteasome, leaving low levels of intact GFP fluorescence (Figure 4b). Strikingly, treatment of cells with rapaprotin led to significantly increased GFP accumulation, similar to bortezomib (Figure 4b). Rapaprotin inhibited proteasome activity in FZ120-H929 cells with an IC_50_ of 817 nM (Figure 4c) at 8 hours according to flow cytometry analysis, reasonably higher than its IC_50_ of 174.8 nM (Figure S2d) in a 72-hour resazurin viability assay due to difference in treatment time. In contrast, rapaprotin failed to cause GFP accumulation in HEK293T cells expressing the mDHFR-GFP reporter, consistent with the insensitivity of this cell line to rapaprotin (Figure S1g).

**Figure 4.**
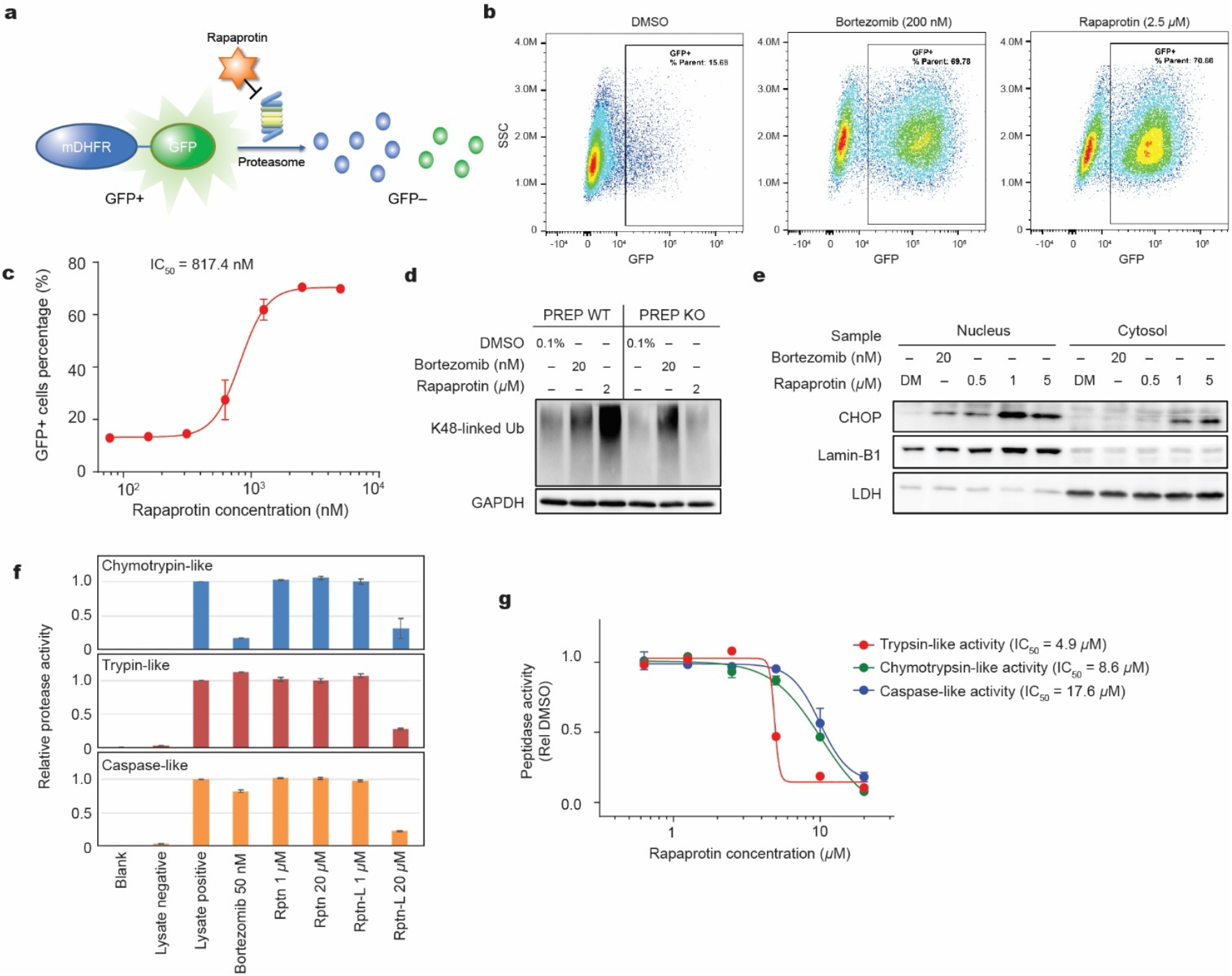
Rapaprotin inhibits ubiquitin-proteasome pathway (UPP) and induces unfolded-protein response (UPR) in NCI-H929 cells, and its activated form, rapaprotin-L, inhibits proteasome activity in vitro. (a) The proteasome reporter cell FZ120-H929 bears a destabilized mutant DHFR tethered to GFP, causing constant degradation of GFP in NCI-H929 cells, inhibition of proteasome leads to increase of GFP+ population. (b) Bortezomib or rapaprotin treatment in proteasome reporter cell line resulted in a large GFP+ population, indicating proteasome inhibition. (c) Dose-dependent inhibition of proteasome by 8-hour treatment with rapaprotin determined by flow cytometry. (d) Western blot shows rapaprotin treatment caused dose-dependent accumulation of K48-linked Ub proteins in WT NCI-H929, but not in PREP KO cells. (e) Western blot shows dose-dependent CHOP activation by rapaprotin in the nucleus and cytosol after 8-hour treatment with rapaprotin. (f) Rapaprotin-L inhibits all three activities of proteasome from the lysate of NCI-H929, while rapaprotin does not. (g) Dose dependent inhibition of proteasome activity by rapaprotin-L in the lysate of NCI-H929. Please refer to Supporting Information for uncropped blots for Figure 4d and e.

Inhibition of the ubiquitin-proteasome pathway in MM cells is known to cause a number of phenotypic changes, including accumulation of polyubiquitinated proteins, induction of PARP cleavage and activation of the unfolded protein response (UPR) pathway. Rapaprotin induced accumulation of K48-linked polyubiquitinated proteins in NCI-H929 cells, but not in PREP-KO cells, demonstrating different selectivity between rapaprotin and bortezomib (Figure 4d). As expected, rapaprotin also induced a nuclear accumulation of CHOP, a marker of the UPR pathway (Figure 4e). CHOP, as well as the proteasome substrates β-catenin (*26*) and p53, were upregulated by rapaprotin in a dose- and time-dependent fashion (*25, 27*) (Figure 4e, S4a, b). Activation of IRE1-α was also observed, indicating the onset of UPR (Figure S4c).

### Rapaprotin-L, but not the cyclic rapaprotin, inhibits protease activities of the proteasome

The inhibition of the mDFHR-GFP reporter by rapaprotin raised the possibility that rapaprotin, and more likely, rapaprotin-L, may directly affect the proteasome. We determined whether rapaprotin inhibited the proteasome. Unlike bortezomib, however, rapaprotin had no effect on the protease activity of any of the three active β-subunits in cell-free assays utilizing NCI-H929 cell lysate (Figure 4f), consistent with the notion that rapaprotin is a latent and inactive pro-drug. In contrast to rapaprotin, the synthetic rapaprotin-L inhibited all three β-subunits at 20 µM in NCI-H929 cell lysates (Figure 4f) while bortezomib selectively inhibited only the chymotrypsin-like activity of the β5 subunit. Among the three active b subunits, rapaprotin-L appeared slightly more potent against the trypsin-(IC_50_ = 4.9 µM) and chymotrypsin-like (IC_50_ = 8.6 µM) activities than the caspase-like activity (IC_50_ = 17.6 µM) (Figure 4g). We note that these differences in IC_50_ values are quite small and can be attributed to the different specificity to the proteasome and rates of cleavage of each of the probes. Despite the relatively high IC_50_ values, the accumulated intracellular concentration of rapaprotin-L (over 120 µM after 8 h incubation, Figure S2c) is more than sufficient to inhibit the activities of all three β-subunits in a cellular context. To further verify the effects of rapaprotin-L on the proteasome and explore the subunit targeting specificity, we determined its effects on purified human 26S proteasome. Similar inhibitory activity of rapaprotin-L was observed with the purified proteasomes (Figure S5).

### Rapaprotin-L induces 26S proteasome disassembly

To investigate the kinetics of the pharmacological action of rapaprotin-L on the proteasome, we conducted time-resolved cryo-EM analysis of the human 26S proteasome treated with rapaprotin-L. Specifically, we imaged the cryo-EM samples flash-frozen on the purified human 26S proteasome that was incubated with rapaprotin-L for 15 and 90 minutes, respectively, and collected a large dataset on the cryo-EM samples for each condition (Table S2). Using a deep-learning-enhanced 3D classification method recently developed (*28*), we were able to classify and identify proteasome particles of very low abundance of both distinct conformations and compositions (Figure S6).

During substrate degradation, the 26S proteasome undergoes a series of conformational changes, typically consisting of ubiquitin recognition in state E_A_, substrate deubiquitylation by RPN11 in state E_B_, initiation of substrate translation in state E_C_ and processive degradation in state E_D_ (*29*). Surprisingly, we identified two novel intermediate proteasome assembles with the complete absence of the lid subcomplex, whereas the complete module of heterohexameric AAA-ATPase motor is attached to the 20S core particle (Figure 5a, Figure S7). One intermediate complex exhibits the density of RPN1 that is attached to the AAA-ATPase in a conformation closely resembling that of a substrate-engaged 26S proteasome in state E_D2_ conformation (*29*). We name this as ‘RPN1-ATPase-CP complex’. In contrast, the RPN1 density is entirely absent in the other intermediate assembly that is referred to as ‘ATPase-CP complex’ and is virtually identical to the ATPase and CP conformations in the 26S proteasome in the resting state (S_A_) (Figure 5c) (*30*). At 15 min of rapaprotin-L treatment, only the ATPase-CP but not RPN1-ATPase-CP complex, which is only v~5% in abundance among all imaged proteasome particles, was found by in-depth 3D classification using AlphaCryo4D and refined to 3.8-Å resolution (Figure 5b). By contrast, at 90 min after addition of rapaprotin-L, both RPN1-ATPase-CP and ATPase-CP complexes were discovered by AlphaCryo4D (*28*) classification and were refined to 4.1 and 3.1 Å, respectively (Figure S7). The cryo-EM maps of ATPase-CP complex separately reconstructed from the datasets from the 15- and 90-min rapaprotin-L treatment appear to be virtually identical, confirming that it represents a stable disassembly intermediate of the 26S proteasome. In contrast to the lack of observation of free 20S CP under the condition of 15 min rapaprotin-L treatment, the majority of proteasome particles was disassembled into free 20S CP (54.0%) and ATPase-CP complex (25.9%) 90 min after rapaprotin-L treatment (Figure 5b). These observations suggest that rapaproin-L may manifest its inhibitory effects on the proteasome through induction of the dissociation of the 19S RP from the CP complex.

**Figure 5.**
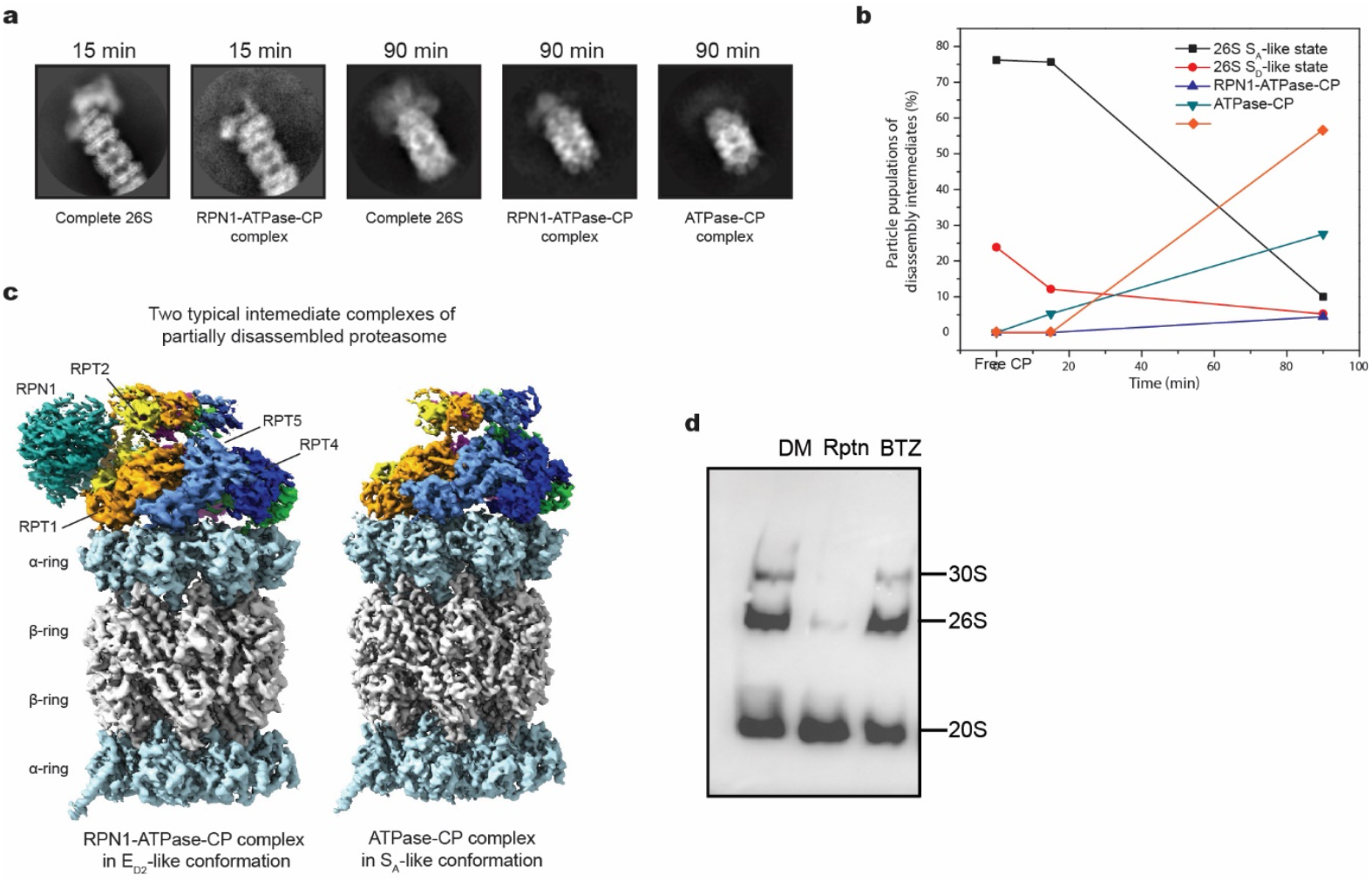
Time-resolved cryo-EM analysis of rapaprotin-L-induced human 26S proteasome disassembly intermediates. (a) Typical 2D class averages of 26S disassembly intermediates as compared to that of the complete 26S assembly, from cryo-EM datasets imaged at 15 and 90 min after rapaprotin-L addition to the purified human 26S proteasome. The lack of the density of lid subcomplex is unambiguously visible. (b) Time-resolved populations of 26S disassembly intermediates show that there are two major intermediate complexes of partially disassembled proteasome, i.e., one that is composed of RPN1, AAA-ATPase hexameric motor and CP, and the other that is composed of only AAA-ATPase and CP. The data points for 0 min was taken from the published results in ref. 32. (c) Cryo-EM reconstructions of two typical intermediate complexes reconstructed from the dataset collected at 90 min after the mixture of rapaprotin with human 26S proteasome. (d) Rapaprotin causes the disassembly of both 26S and 30S proteasome in cells. NCI-H929 cells were incubated with vehicle (0.1% DMSO or DM), Rapaprotin (Rptn), and Bortezomib (BTZ) for 8 h. Cells were harvested and lysed; cell lysates were subjected to 4% native polyacrylamide gel electrophoresis followed by transfer onto a PVDF membrane. The membrane was blotted using antibodies against the β5 subunit of 20S core particle. Please refer to Supporting Information for uncropped blots for Figure 5d.

To validate the effect of rapaprotin-L on the 26S proteasome observed in vitro, we determined the effect of rapaprotin on the proteasome assembly in cells. Thus, NCI-H929 cells were treated with DMSO (control), 1 µM rapaprotin and 4 nM bortezomib, respectively, for 8 h. The cells were harvested and cell lysates were subjected to native gel electrophoresis followed by Western blot analysis using polyclonal antibodies against the b5 subunit of the 20S CP. In the absence of rapaprotin, both 26S and 20S proteasomes are abundant with a small amount of 30S present (Figure 5d). Treatment with rapaprotin led to the disappearance of both the 26S and 30S species. In contrast, bortezomib had negligeable effect on the relative abundance of 26S and 30S. These results indicate that rapaprotin causes the 26S disassembly in multiple myeloma cells, corroborating the observations made with 26S proteasome using cryo-EM in vitro.

To understand if rapaprotin-L inhibits the proteasome by directly attacking the active site of the CP, we examined all active sites in the three proteolytic β-subunits of the proteasome in all our cryo-EM reconstructions. No additional densities were observed in all active sites in the CP, suggesting that rapaprotin-L does not bind to the active site grooves of the CP. Rather, it appeared to destabilizes the 19S RP, causing its dissociation from the 20S CP. In eukaryotes, the 19S RP is required for the 26S proteasome activation for ubiquitin-mediated substrate degradation. Rapaprotin-L-induced disassembly of RP from the CP deactivates the proteasome, therefore severely impairing proteasome function for degrading ubiquitin-tagged substrates. This observation explains why rapaprotin-L can inhibit all three proteolytic activities of the 26S proteasome together. In this regard, rapaprotin-L is strongly reminiscent of intracellular proteasome-interacting protein Ecm29, which is known to induce the disassembly of 26S proteasome, thereby inhibiting proteasome function (*31, 32*). Thus, induction of proteasome disassembly by small-molecule compound appears to be a viable, novel approach to proteasome inhibition.

### Rapaprotin synergizes with bortezomib and re-sensitizes bortezomib-resistant MM cells

Rapaprotin-L causes disassembly of the 26S proteasome, impairing the processing and degradation of polyubiquitinated substrates in a mechanism distinct and orthogonal to that of existing CP inhibitors, raising the possibility that rapaprotin may be synergistic with them for the induction of apoptosis in MM cells. We thus determined full dose-response curves for rapaprotin in the absence and the presence of increasing concentrations of bortezomib (Figure 6a). As the concentrations of bortezomib were increased, there was a corresponding decrease in the IC_50_ value for rapaprotin. The ZIP analysis revealed that there is strong synergy between rapaprotin and bortezomib with an average synergy score of 15.5 (Figure 6b). Intriguingly, the synergy between rapaprotin and bortezomib was observed in NCI-H929 cells but not the primary HUVEC (Figure 6c). The synergy between rapaprotin and bortezomib suggests that they may be used in combination to achieve greater anticancer efficacy. As expected, we also found that rapaprotin is synergistic with carfilzomib (CFZ, Figure S8a) and ixazomib (IXZ, Figure S8b).

**Figure 6.**
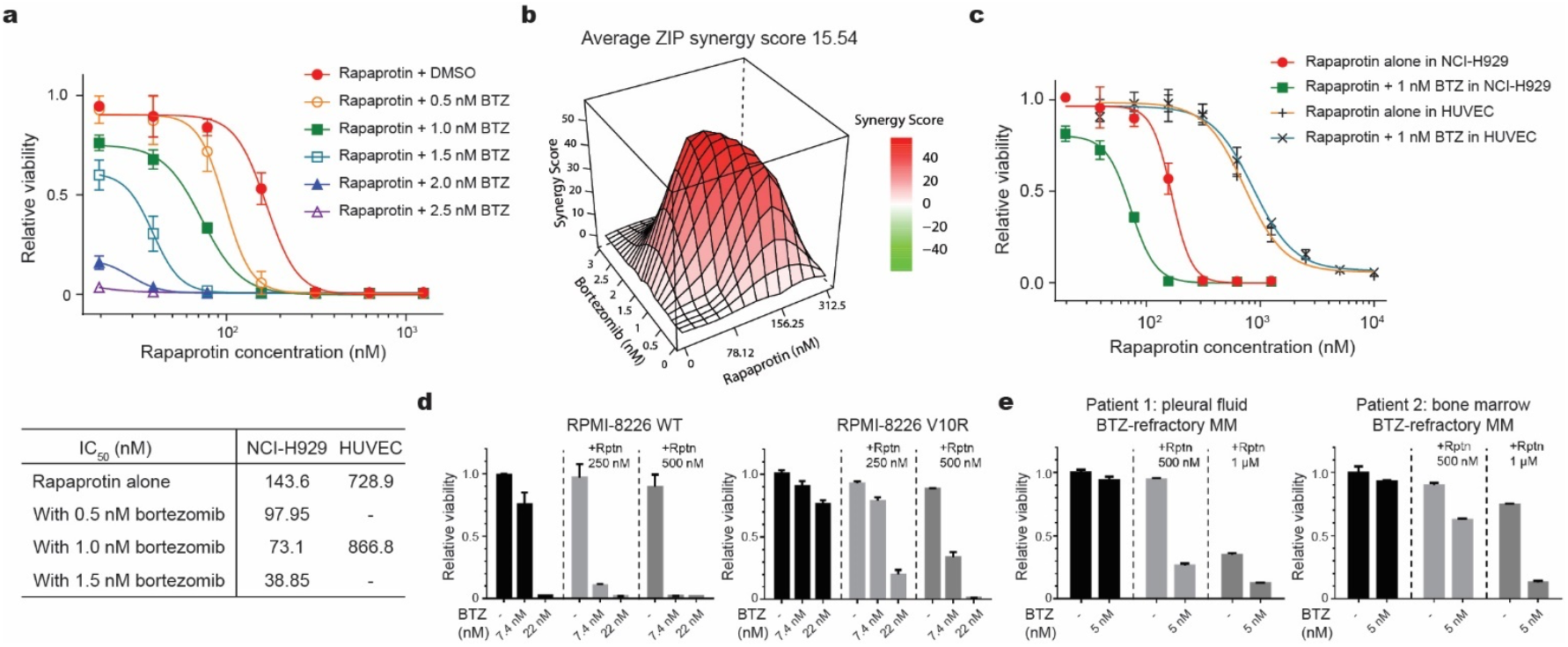
Rapaprotin exhibits synergy with FDA-approved proteasome inhibitor bortezomib. (a) Synergistic inhibition of rapaprotin and bortezomib in HCI-H929. (b) Zero interaction potency (ZIP) synergy plot of 72-hour treatment of NCI-H929 cells with rapaprotin in combination with bortezomib. Synergy analysis and results were generated with SynergyFinder. Positive scores indicate synergy, and negative scores indicate antagonism. (c) Combination treatment of rapaprotin and Bortezomib in NCI-H929 and HUVEC cells. (d) Wild-type (WT) and bortezomib-resistant (V10R) MM cell lines were treated with Bortezomib and rapaprotin for 48 hours followed by MTT assay. (e) CD138+ MM patient samples were treated as in (d). Rptn, rapaprotin; BTZ, bortezomib.

Since drug resistance remains the major clinical challenge for patients treated with bortezomib and other existing proteasome inhibitor drugs, we next sought to determine whether rapaprotin can overcome such resistance. To accomplish this, we first generated dose-response curves to rapaprotin, bortezomib, or the combination in wild-type RPMI-8226 compared to the bortezomib-resistant cell line model RPMI-8226 V10R. Exposing either wild-type or V10R bortezomib-resistant cells to 0.25 or 0.5 µM dramatically enhanced their sensitivity to bortezomib (Figure 6d). We also examined the combination of bortezomib with E27-9 in MM primary samples derived from two bortezomib-refractory patients. Similar to RPMI-8226 V10R, the bortezomib-refractory MM cells from both patients were inhibited by nearly 90% in the presence of the combination of bortezomib and E27-9, but not either alone (Figure 6e). These results strongly suggest that rapaprotin is capable of sensitizing resistant MM cells to bortezomib.

The proteasome is one of the most complex molecular machineries that possess many enzymatic activities requiring remarkable allosteric coordination, dynamic timing and bidirectional regulation (*33, 34*). In cells, there are two categories of natural inhibitors of the proteasome. One is the PI31 protein that can enter the CP chamber and directly binds all six proteolytic active sites (*35, 36*). The other is the Ecm29 protein that was believed to instead induce the RP-CP disassembly to quench the 26S proteasome activity (*31, 32*). Those well-known proteasome inhibitors, such as bortezomib, carfilzomib and ixazomib, are similar to PI31 by directly binding the proteolytic sites of active β-subunits to block substrate degradation. In contrast, rapaprotin-L surprisingly bears mechanistic similarity to Ecm29 for inducing the disassembly of the 26S proteasome.

The precise binding site for rapaprotin-L remains elusive. Despite our exhaustive analysis of the cryo-EM datasets, we did not detect any complexes between rapaprotin-L and the proteasome subunits both prior to and after its disassembly. Coincidently, rapamycin and its linearized analog were reported to inhibit the 20S proteosome activity and block binding of 19S to the 20S core particle at relatively high concentrations in vitro (*37*). More recently, a simplified pipecolic ester derivative derived from the FKBD of rapamycin was shown to inhibit the 20S proteasome activity. It was hypothesized that rapamycin binds to the a face of the 20S CP and molecular docking suggested that the pipecolate-based inhibitor binds to the interface between the a2 and a3 subunits on the 20S CP (*38*). Although rapaprotin bears structural similarity to the FKBD domain of rapamycin and the pipecolic ester, it differs from them in several ways. First, cyclic rapaprotin is in active and required cleavage by PREP to be active. Second, cyclic rapaprotin has no effect on the protease activity of 20S and rapaprotin-L inhibits all three protease activities of the 20S. In contrast, the cyclic rapamycin and the pipecolate ester selectively inhibit two of the three protease activities (b1 and b5) of 20S while stimulating (rapamycin) or sparing (pipecolate ester) the trypsin-like (b2) activity. Third, all inactive rapaprotin analogs share the same pipecolate-containing FKBD as rapaprotin, ruling out the possibility that the similar FKBDs in these molecules are responsible for their effects on the proteasome. Future work will be needed to determine the precise binding site of rapaprotin-L on the 26S proteasome and its relationship to the putative rapamycin- and pipecolic ester-binding sites. To our knowledge, a latent inhibitor selectively activated to induce disassembly of a multiprotein complex to impair the target function presents a new inhibition/intervention paradigm in the mode of action by a small molecule. Proteasome inhibition constitutes a major therapeutic option for MM and lymphoma patients. The three FDA-approved proteasome inhibitors do have major limitations including toxicity and drug-resistance. Recent studies on the mechanism of drug resistance of proteasome inhibitors have revealed the importance to concurrently blocking more than one of the three active β-subunits of the proteasome. Identification of a proteasome assembly inhibitor, rapaprotin, which exhibits highly selective cellular toxicity in MM cells over primary cells, presents a novel mode of proteasome inhibition by inducing the 26S proteasome disassembly. Upon intracellular activation by the cellular protease PREP, rapaprotin-L that bears a negative charge is trapped inside cells, accumulating to an extremely high concentration, causing indirect inhibition of all three proteolytic activities of the proteasome and leading to apoptosis of MM cells.

Rapaprotin has a unique and unprecedented mode of action in comparison with rapamycin and FK506. In particular, the major structural transformation from an inactive cyclic form to an active linear form upon cleavage by PREP has not been seen in other small molecule modulators of proteins. The rapafucin library from which rapaprotin originated was inspired by the unique mode of action of rapamycin, FK506 and cyclosporin A as molecular glues (*39, 40*). Indeed, a new molecular glue from the rapafucin library, a potent and isoform-specific inhibitor of equilibrative nucleoside transporter-1 (ENT1) called rapadocin, has been identified (*20*). Rapaprotin, however, represents a notable departure from molecular glues in its mode of action. Instead, it serves as a molecular transformer. Its inactive cyclic form not only masks its intrinsic biological activity towards the proteasome, but also allows it to penetrate the plasma membrane to gain cellular entry. Once inside cells, PREP-mediated cleavage transforms the cyclic rapaprotin into the linear rapaprotin-L to act on the 26S proteasome. It will be interesting to see if more molecular transformers like rapaprotin may emerge from the rapafucin library.

It was unexpected that rapaprotin is activated by the intracellular protease PREP to become an active proteasome inhibitor. PREP enzyme is widely expressed in different types of cells and their expression level can vary among different cancer cells (*41-44*). A major determinant for PREP substrates is a proline residue at the P1 position. Moreover, the natural substrates for PREP are linear proteins or peptides. The FKBD moiety of both rapamycin and FK506 contain a pipecolate lactone that mimics a proline residue, similar to rapaprotin. But they are not subjected to PREP cleavage. Surprisingly, the same pipecolate lactone in rapaprotin became susceptible to PREP-catalyzed hydrolysis. A comparison between the FKBDs in rapamycin/FK506 and that in rapaprotin revealed that the substituted cyclohexyl sidechain in rapamycin that corresponds to the sidechain of the P1’ position of PREP substrates is replaced by a dimethoxyphenyl sidechain, which is likely responsible for the fortuitous gain in sensitivity to PREP. Indeed, we found that modification of the dimethoxyphenyl moiety in rapaprotin, i.e., removal of one of both methoxy groups or conversion of a methoxy group into a hydroxyl group, renders the corresponding rapaprotin analogs resistant to PREP, abolishing the proapoptotic activity in MM cells. The unique rapafucin scaffold may serve as a novel delivery vehicle for other drugs by swapping them with the effector domain of rapaprotin.

It is noteworthy that although intracellular concentration of cyclic rapaprotin was only ~7 µM, the concentration of rapaprotin-L was close to 70 µM three hours after treatment with rapaprotin, and over 120 µM eight hours post rapaprotin treatment (Figure S2C), likely due to lack of cell membrane permeability with its newly generated carboxylate group. For the same reason, extracellular rapaprotin-L can hardly enter the cells, as evidenced by lack of cytotoxicity of rapaprotin-L at concentrations up to 5 µM (Figure S8c). The low membrane permeability is responsible for trapping rapaprotin-L inside the cells, assuring high local concentrations of the active species. The time-dependent accumulation of rapaprotin-L inside cells also offers an explanation for the moderate potencies of rapaprotin-L for inhibition of the proteolytic activity of the proteasome in vitro (Figure 4g) and the high potency of rapaprotin in MM cells (Figure 1c).

The selectivity of rapaprotin for cancer cells over normal cells in contrast to bortezomib and carfilzomib can be attributed to at least two factors. One is the inherent sensitivity of the target cell to proteasome inhibition based on factors such as proteasome load/capacity ratio (*45, 46*), which is a pattern observed with other proteasome inhibitors. For example, the heaviest loaded NCI-H929 is also the most sensitive cell line to rapaprotin, while lighter loaded cells, such as KMS-12 and RPMI-8226, are less sensitive (Figure S8d). The second factor is an added layer of selectivity which comes from its property as a pro-drug that is activated by an endogenous endopeptidase. Although we did not identify any strict correlation between PREP mRNA expression and rapaprotin sensitivity in the cell line encyclopedia assay, small differences in PREP activity between cell types could lead to selective accumulation over time. The logarithm of IC_50_ shows a negative correlation with the intracellular rapaprotin-L concentration (Figure S8e), underscoring the contribution of PREP activity to the effectiveness of rapaprotin.

## Conclusion

In this work, we identified rapaprotin as a novel inhibitor of the 26S proteasome with a unique mechanism of action. The cyclic rapaprotin is a latent pro-drug that can penetrate plasma membrane of cancer cells. Upon cellular entry, it is cleaved by the cytosolic protease PREP to become rapaprotin-L that inhibits the 26S proteasome. Unlike existing proteasome inhibitors used in the clinic that target the active b subunits of the 20S CP, rapaprotin-L induces the disassembly of the 26S proteasome, leading to the eventual dissociation of the 19S RP from the 20S CP. This selective inhibition of the 26S proteasome function confer several advantages to rapaprotin. Rapaprotin exhibited greater selectivity for MM cells over normal endothelial cells, suggesting that it may have lower toxicity than existing proteasome inhibitors. It also showed dramatic synergy with bortezomib and is capable of sensitizing drug-resistant MM cells from patients to bortezomib, making it possible to use bortezomib-rapaprotin combination with a lower dose of bortezomib, both reducing the toxicity of bortezomib and enabling a more durable response for MM patients. Thus, rapaprotin not only serves as a novel chemical probe for studying proteasome dynamics and function, but also holds promise as a first-in-class therapeutic for diseases driven by dysregulated proteasome activity.

## Supporting information

Figures S1-9, Tables S1-2, Materials and Methods, Chemical Synthesis

## Acknowledgements

This work was supported in part by NIH (R01GM145793) and the Commonwealth Fund (CBG & JOL), FAMRI (JOL), a Damon Runyon Postdoctoral Fellowship (HP), and grants from the National Natural Science Foundation of China (12125401, 12090054 and 11774012) and the Beijing Natural Science Foundation (Z180016/Z18J008) (YM). The cryo-EM data were collected at the Cryo-EM Core Facility Platform and Laboratory of Electron Microscopy at Peking University. The data processing was performed in the High-Performance Computing Platform, the Weiming No. 1 and Life Science No. 1 supercomputing systems at Peking University. We thank C. Yang for discussions related to CRISPR screening, and Z. Guo, N. Li for assistance in cryo-EM sample screening. We are grateful to Dr. Philip Cole for critical comments on the manuscript.

