## Supplementary material for "Rapaprotin is Activated by an Endopeptidase to Disassemble 26S Proteasome": Figures S1-9, Tables S1-2, Materials and Methods, Chemical Synthesis

**This PDF file includes**

Figures S1 to S9

Tables S1 to S2

Materials and Methods

Chemical Synthesis

SI References

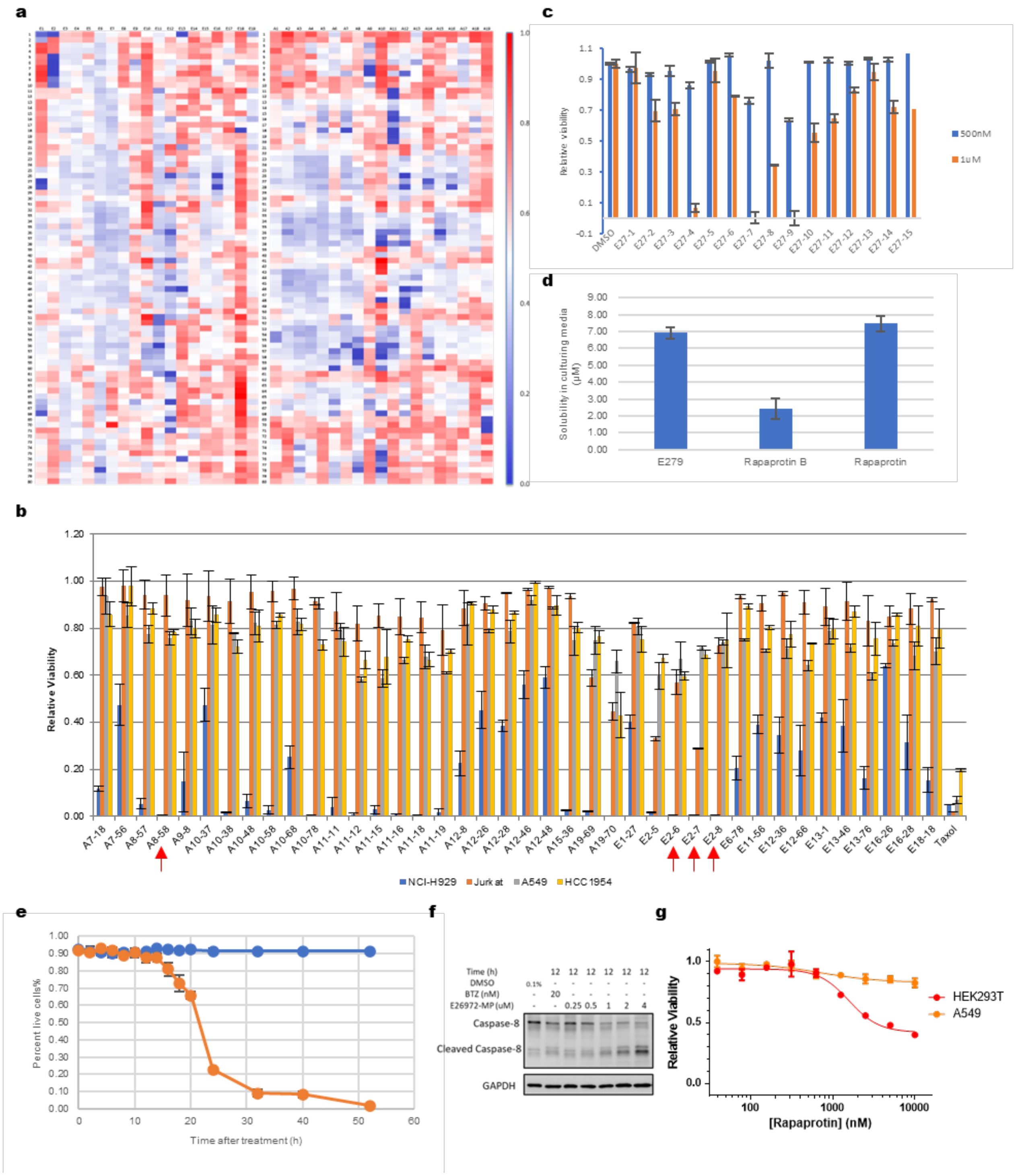

**Fig. S1**. **Screen and discovery of paraprotin as a proapoptotic agent**. (**a**) Heatmap of the outcome of the screening of rapafucin libraries using resazurin viability assay in NCI-H929 cells. FKBD11- (left) and FKBD10-containing libraries (right) were screened at a final concentration of 200 nM per compound or 3 μM total rapafucin in each well for 72 hours prior to analysis. Color scale shows remaining viability from 0.0 (100% inhibition) to 1.0 (no inhibition). The arrow points to one of the most potent pool of hits at Row 6,

Column E2, that was further pursued. (**b**) Bar graph of re-screening of 40 top hits using resazurin viability assay in NCI-H929, Jurkat, A549, and HCC1954. Cells were treated with pools of rapafucins at a concentration of 200 nM per compound, or 3μM total rapafucin in each well for 72h prior to analysis. The graph is a representative of 3 independent experiments. Error bars represent s.e.m.; data are mean ± s.e.m. (**c**) Bar graph of re-screening of 15 decoded individual compounds in rapafucin pooled hit E27 using resazurin viability assay in NCI-H929. E27-9 was found to be the most active compound among these 15 rapafucins. Cells were treated with individual compounds at a concentration of 500 nM and 1μM in each well for 72 h prior to analysis. The graph is a representative of 3 replicates. Error bars represent s.e.m.; data are mean ± s.e.m. (**d**) Bar graph showing solubility of E27-9, rapaprotin B, and rapaprotin in cell culturing media (RPMI 1640 with 10% FBS) as determined with HPLC-MS. The graph is a representative of 3 replicates. Error bars represent s.e.m.; data are mean ± s.e.m. (**e**) Cell counting with trypan blue staining in NCI-H929 cells with treatment of rapaprotin at 1 μM. (**f**) Western blot shows concentration dependent activation of Caspase 8 after treatment with rapaproptin. (**g**) Dose-dependent cytotoxicity of rapaprotin in HEK293T ($EC_{50}$ 1575 nM) and A549 ($IC_{50}$ >10000 nM) cells. Cell viability was measured using resazurin viability assay. Error bars represent s.d.; data are mean ± s.d; n = 3 independent experiments.

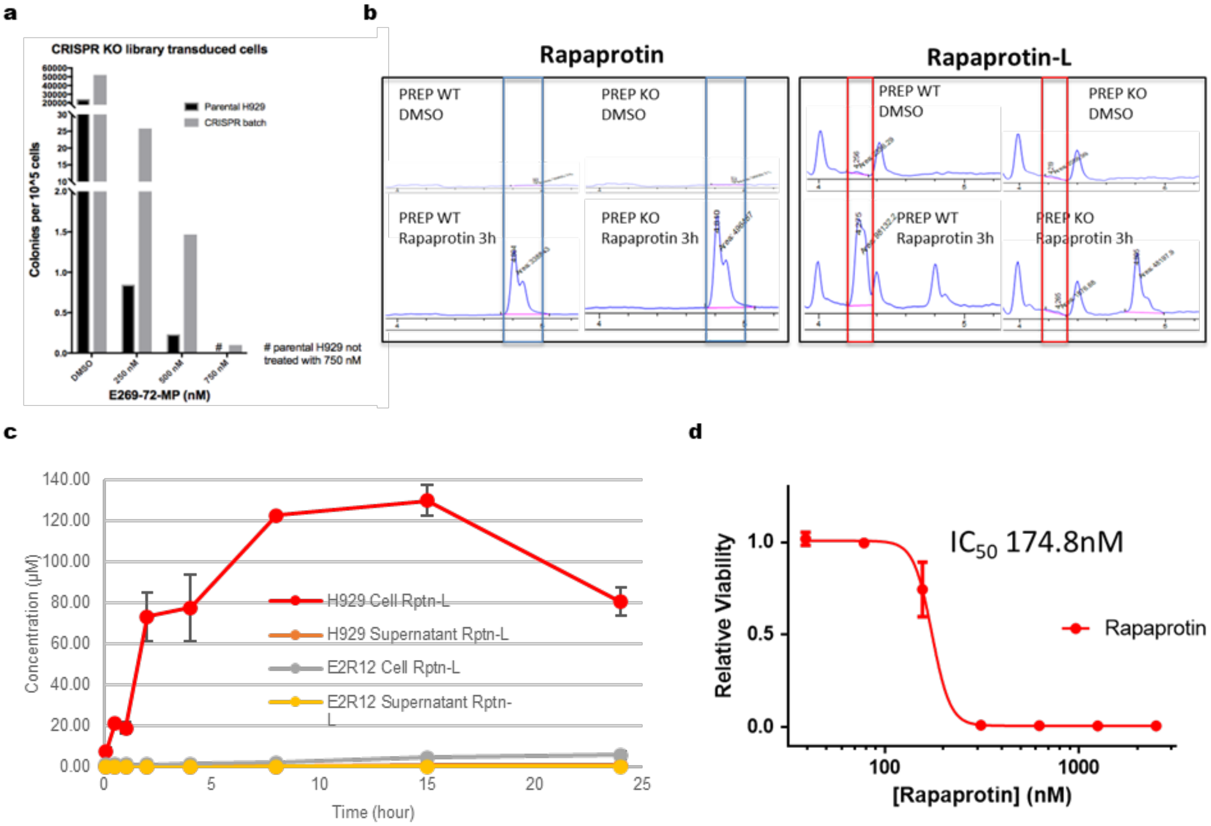

**Fig. S2. Evidence for the activation of rapaprotin by prolyl endopeptidase and the action of rapaprotin-L**. (**a**) Determining concentration of rapaprotin used for CRISPR screening. (**b**) HPLC detection chromatogram of rapaprotin and its cleaved product rapaprotin-L in WT NCI-H929 and PREP knock-out strain E2R12 with or without treatment of rapaprotin for 3 hours. rapaprotin shows a peak at 4.8-4.9 min, rapaprotin-L shows a peak at 4.2-4.3 min. LC-MS was run with a Selective Ion Monitoring (SIM) mode, and chromatogram show ion extraction result for m/z 1334.6 (Rapaprotin+$H^+$) to represent rapaprotin, and m/z 1352.6 (rapaprotin+$H_2O$+$H^+$) to represent rapaprotin-L. (**c**) Measurement of intracellular and extracellular rapaprotin and rapaprotin-L upon treatment in PREP-WT NCI-H929 cells and PREP-KO NCI-H929 cells (E2R12). (**d**) Dose-dependent cytotoxicity of rapaprotin in proteasome reporter cell line FZ120-H929. Cell viability was measured using resazurin viability assay. Error bars represent s.d.; data are mean ± s.d; n=3 independent experiments.

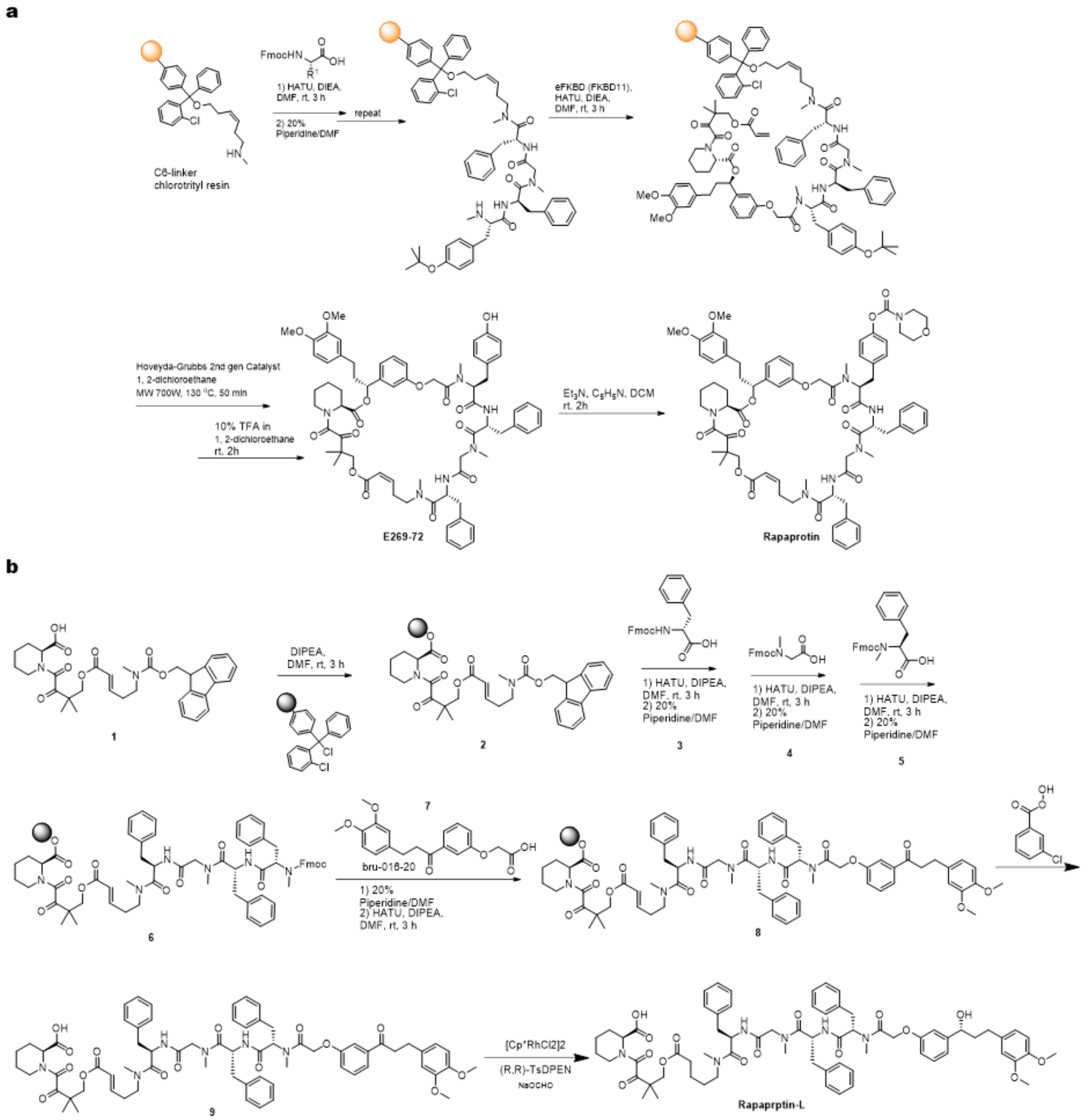

**Fig. S3. Synthesis** of rapaprotin and rapaprotin-L**.** (**a**) Synthesis of Rapaprotin through solid phase chemistry. (**b**) Synthesis of Rapaprotin-L through solid phase chemistry.

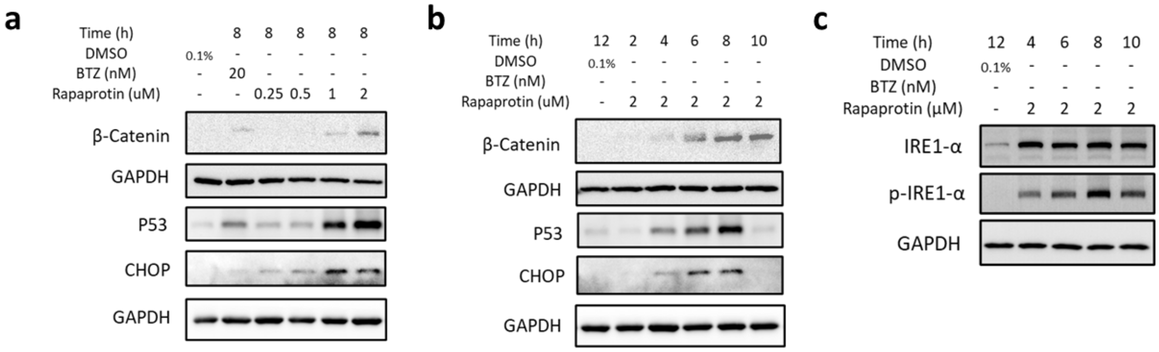

**Fig. S4.** (**a, b**) Western blot shows dose-dependent and time-dependent up-regulation of proteasome substrates β-Catenin and P53, and activation of CHOP by rapaprotin in NCI-H929. (**c**) Western blot shows time dependent up-regulation and phosphorylation of IRE1-α induced by rapaprotin in NCI-H929.

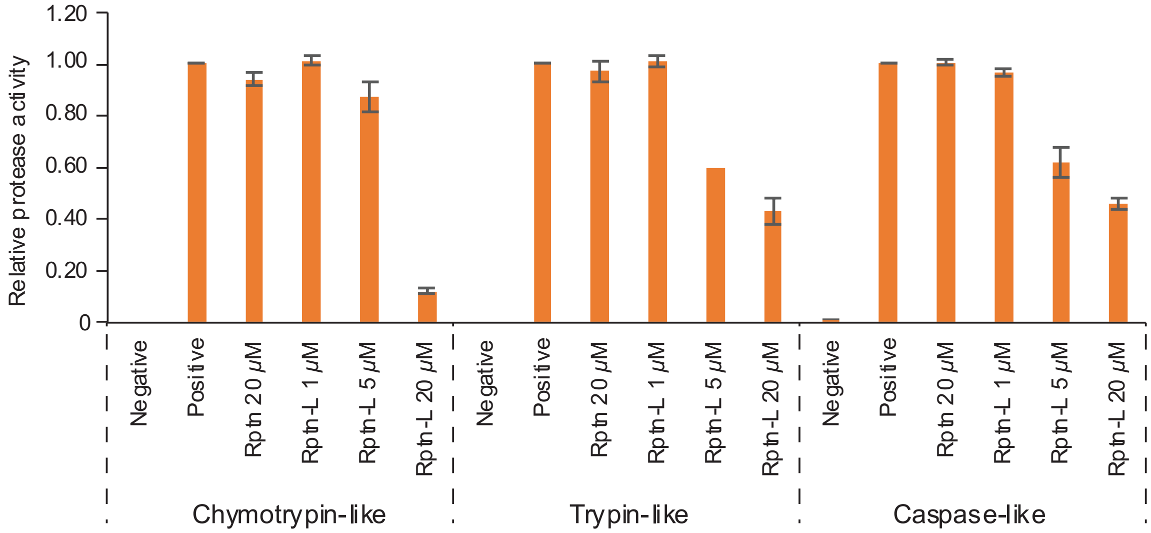

**Fig. S5.** Rapaprotin-L inhibits the activity of recombinant 26S proteasome. BTZ, bortezomib.

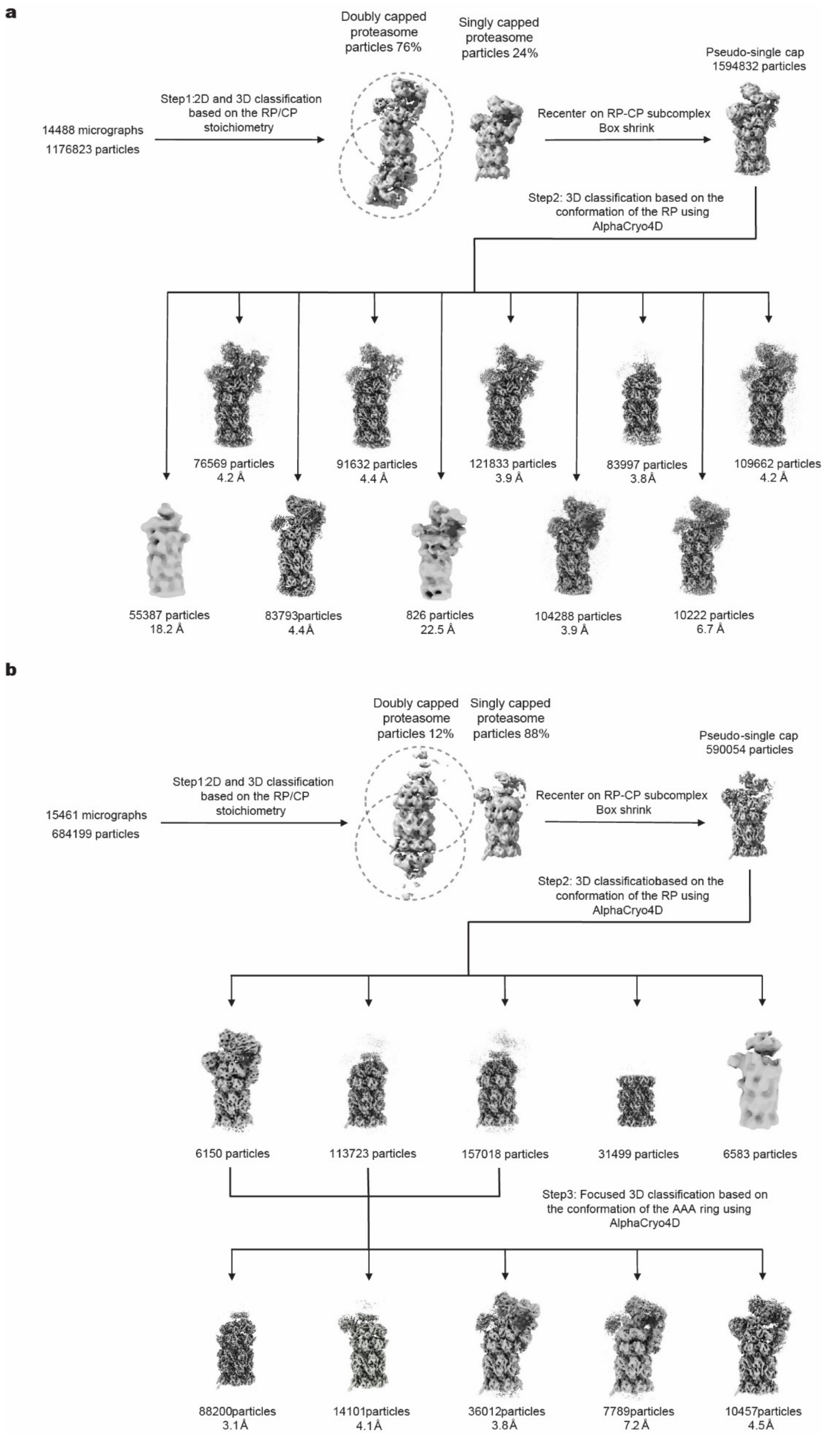
a
Doubly capped proteasome particles 76%
Singly capped proteasome particles 24%
Pseudo-single cap 1594832 particles
Step1:2D and 3D classification based on the RP/CP stoichiometry
14488 micrographs
1176823 particles
Recenter on RP-CP subcomplex
Box shrink
Step2: 3D classification based on the conformation of the RP using AlphaCryo4D
76569 particles
4.2 Å
91632 particles
4.4 Å
121833 particles
3.9 Å
83997 particles
3.8Å
109662 particles
4.2 Å
55387 particles
18.2 Å
83793particles
4.4Å
826 particles
22.5 Å
104288 particles
3.9 Å
10222 particles
6.7 Å
b
Doubly capped proteasome particles 12%
Singly capped proteasome particles 88%
Pseudo-single cap 590054 particles
Step1:2D and 3D classification based on the RP/CP stoichiometry
15461 micrographs
684199 particles
Recenter on RP-CP subcomplex
Box shrink
Step2: 3D classificatiobased on the conformation of the RP using AlphaCryo4D
6150 particles
113723 particles
157018 particles
31499 particles
6583 particles
Step3: Focused 3D classification based on the conformation of the AAA ring using AlphaCryo4D
88200particles
3.1Å
14101particles
4.1Å
36012particles
3.8Å
7789particles
7.2Å
10457particles
4.5Å

**Fig. S6. Cryo-EM data processing workflow**. The workflow diagram illustrates the major steps of our focused 3D classification strategy using AlphaCryo4D for the dataset collected from the 26S human proteasome sample flash-frozen 15 min (**a**) and 90 min (**b**) after mixture of rapaprotin with the proteasome.

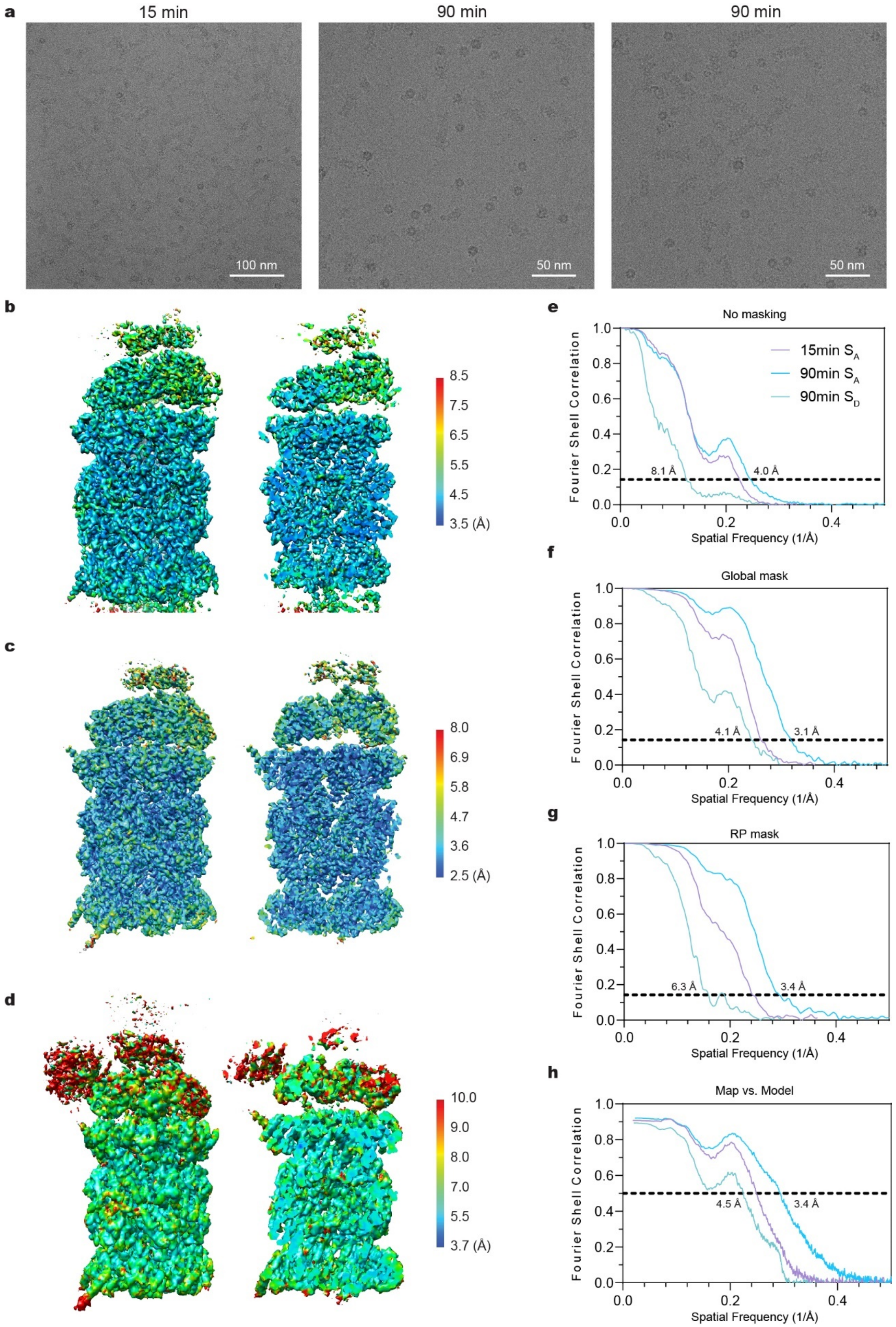
a
15 min
90 min
90 min
100 nm
50 nm
50 nm
b
8.5
7.5
6.5
5.5
4.5
3.5 (Å)
c
8.0
6.9
5.8
4.7
3.6
2.5 (Å)
d
10.0
9.0
8.0
7.0
5.5
3.7 (Å)
e
No masking
15min S_A
90min S_A
90min S_D
8.1 Å
4.0 Å
Fourier Shell Correlation
Spatial Frequency (1/Å)
f
Global mask
4.1 Å
3.1 Å
g
RP mask
6.3 Å
3.4 Å
h
Map vs. Model
4.5 Å
3.4 Å

**Fig. S7. Cryo-EM reconstructions and resolution measurement.** (**a**-**b**) Local resolution estimation of the cryo-EM reconstructions of the ATPase-CP intermediate assembles from the dataset collected at 15 min (**a**) and 90 min (**b**) rapaprotin addition and the RPN1-ATPase-CP intermediate assembly (**c**) from the dataset collected at 90 min rapaprotin addition. d,e, Gold-standard Fourier shell correlation (FSC) plots of the three proteasome subassemblies calculated without (**d**) and with (**e**) masking the separately refined half-maps. (**f**) Gold-standard FSC plots of the three proteasome subassemblies calculated with masking the RP component of the separately refined half-maps. (**g**) Model-map FSC plots calculated between each reconstruction and its corresponding atomic model.

a

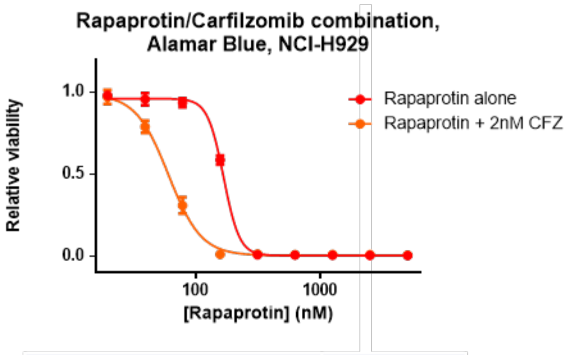

| Drug combination | $IC_{50}$ (nM) |
|---|---|
| Rapaprotin alone | 166.7 |
| Rapaprotin + 2nM Carfilzomib | 60.2 |

b

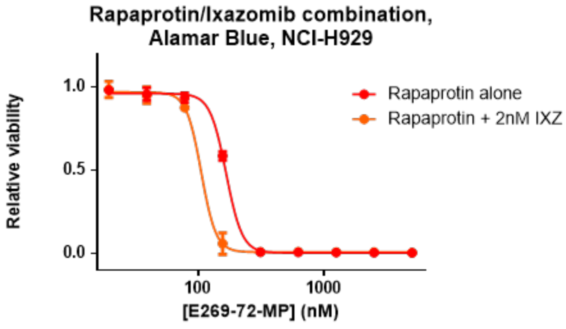

| Drug combination | $IC_{50}$ (nM) |
|---|---|
| Rapaprotin alone | 166.7 |
| Rapaprotin + 2nM Ixazomib | 105.8 |

c

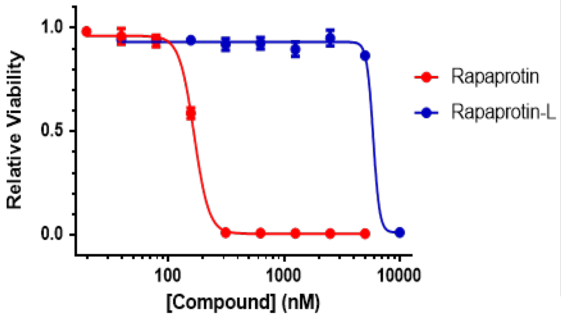

d

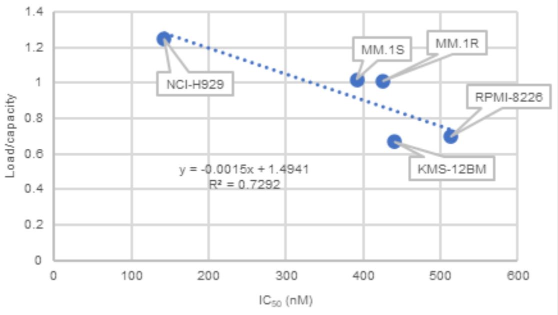

| | $IC_{50}$ (nM) | Load/capacity |
|---|---|---|
| NCI-H929 | 143 | 1.25 |
| MM.1S | 392 | 1.02 |
| MM.1R | 426 | 1.01 |
| KMS-12 | 441 | 0.67 |
| RPMI-8226 | 513 | 0.7 |

e

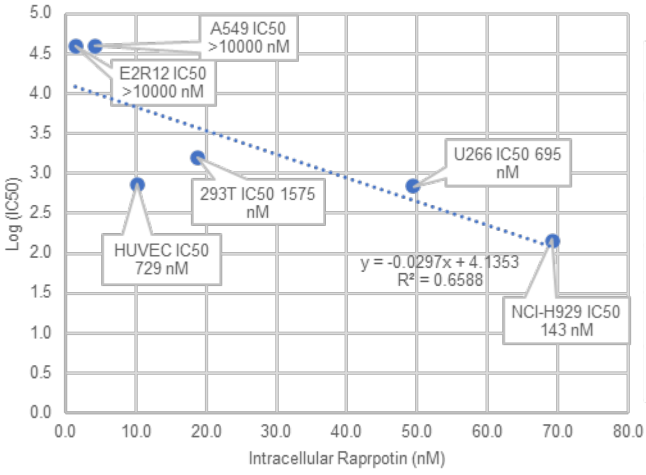

| | Rptn-L (nM) | $IC_{50}$ (nM) | Log $IC_{50}$ |
|---|---|---|---|
| NCI-H929 $IC_{50}$ = 143 nM | 69.2 | 143 | 2.2 |
| U266 $IC_{50}$ = 695 nM | 49.5 | 695 | 2.8 |
| HUVEC $IC_{50}$ = 729 nM | 10.2 | 729 | 2.9 |
| 293T $IC_{50}$ = 1575 nM | 18.8 | 1575 | 3.2 |
| A549 $IC_{50}$ >10000 nM | 4.2 | >10000 | >4.0 |
| E2R12 $IC_{50}$ >10000 nM | 1.4 | >10000 | >4.0 |

**Fig. S8. Synergistic effects of rapaprotin with other proteasome inhibitors.** (**a**) Rapaprotin is synergistic with calfilzomib (CFZ) in NCI-H929 cells. (**b**) Rapaprotin is synergistic with Ixazomib (IXZ) in NCI-H929 cells. (**c**) Dose-dependent cytotoxicity of rapaprotin ($IC_{50}$ 166 nM) and rapaprtin-L ($IC_{50}$ ~5885 nM) in NCI-H929. Cell viability was measured using resazurin viability assay. Error bars represent s.d.; data are mean ±

s.d; n=3 replicates. (**d**) Correlation between the sensitivity of different cell lines (NCI-H929, MM1.S, MM1.R, KMS-12, and RPMI-8226) and their proteasome load/capacity ratios. (**e**) Correlation between the sensitivity of different cell lines (NCI-H929 WT, NCI-H929 PREP-KO, U266, HUVEC, and A549) and the intracellular concentrations of rapaprotin-L.

**Table S1. gRNA sequences and sensitivity to bortezomib and Rapaprotin in resistant cells from CRISPR selection**

| Resistant cell | gRNA | Target protein | $IC_{50}$ of rapaprotin (nM) | $IC_{50}$ of bortezomi (nM) |
|---|---|---|---|---|
| Rptn-r1 (C4A) | CATCGTCAGACAGTATGTTG | PREP | >10000 | 2.9 |
| Rptn-r2 (C4C) | CATCGTCAGACAGTATGTTG | PREP | >10000 | 3.0 |
| Rptn-r3 (HC750-19) | GCAGGAATCCAGTGGCATCG | PREP | >10000 | 3.5 |
| Rptn-r4 (HC750-60) | GAATGTTCTTGACGTCATGG | PREP | >10000 | 3.4 |
| Rptn-r5 (E2R2) | CATCGTCAGACAGTATGTTG | PREP | >10000 | 3.0 |
| Rptn-r6 (E2R3) | GCAGGAATCCAGTGGCATCG | PREP | >10000 | 2.4 |
| Rptn-r7 (E2R11) | GCAGGAATCCAGTGGCATCG | PREP | >10000 | 2.5 |
| Rptn-r8 (E2R12) | GCAGGAATCCAGTGGCATCG | PREP | >10000 | 2.6 |
| Rptn-r9 (E2R21) | TCTTTGTATAAACCTCTGAT | PREP | >10000 | 2.1 |
| Rptn-r10 (E2R37) | GCAGGAATCCAGTGGCATCG | PREP | >10000 | 2.6 |
| Rptn-r11 (E2R45) | GCAGGAATCCAGTGGCATCG | PREP | >10000 | 2.6 |

**Table S2. Cryo-EM data collection, refinement and validation statistics**

| | 15min-c1 (EMD-xxxxx) (PDB XXXX) | 90min-c1 (EMD-xxxxx) (PDB XXXX) | 90min-c3 (EMD-xxxxx) (PDB XXXX) |
|---|---|---|---|
| **Data collection and processing** | | | |
| Magnification | 105,000 | 165,000 | 165,000 |
| Voltage (kV) | 300 | 300 | 300 |
| Electron exposure ($e^{-}/Å^2$) | 50 | 50 | 50 |
| Defocus range (μm) | -0.4 to -5.0 | -0.4 to -5.0 | -0.4 to -5.0 |
| Pixel size (Å) | 0.685 | 0.42 | 0.42 |
| Symmetry imposed | C1 | C1 | C1 |
| Initial particle images (no.) | 1176823 | 684199 | 684199 |
| Final particle images (no.) | 83997 | 88200 | 14101 |
| Map resolution (Å) | 3.8 | 3.1 | 4.1 |
| FSC threshold | 0.143 | 0.143 | 0.143 |
| Map resolution range (Å) | 3.5-8.5 | 2.5-8.0 | 3.7-10.0 |
| **Refinement** | | | |
| Initial model used | | | |
| Model resolution (Å) | 4.1 | 3.4 | 4.3 |
| FSC threshold | 0.5 | 0.5 | 0.5 |
| Model resolution range (Å) | 3.5-8.5 | 2.5-8.0 | 3.7-10.0 |
| Map sharpening *B* factor ($Å^2$) | -65 | 0 | 0 |
| **Model composition** | | | |
| Non-hydrogen atoms | 63361 | 63361 | 68847 |
| Protein residues | 8242 | 8242 | 8979 |
| Ligands | 6 | 6 | 5 |
| *B* factors ($Å^2$) | | | |
| Protein | 181.45 | 119.47 | 167.07 |
| Ligand | 277.82 | 103.44 | 186.53 |
| **R.m.s. deviations** | | | |
| Bond lengths (Å) | 0.004 | 0.004 | 0.006 |
| Bond angles (°) | 0.864 | 0.864 | 1.135 |
| **Validation** | | | |
| MolProbity score | 1.68 | 1.69 | 2.12 |
| Clashscore | 5.34 | 5.65 | 11.05 |
| Rotamers outliers (%) | 0.32 | 0.36 | 0.15 |
| **Ramachandran plot** | | | |
| Favored (%) | 94.16 | 94.37 | 89.66 |
| Allowed (%) | 5.78 | 5.60 | 10.19 |
| Outliers (%) | 0.06 | 0.04 | 0.15 |

### Materials and Methods

#### Cell lines

NCI-H929 (Cat#: CRL-9068), RPMI-8226 (Cat#: CCL-155), U266 (Cat#: TIB-196), Pfeiffer (Cat#: CRL-2632), MM1.S (Cat#: CRL-2974), Jurkat T (Cat#: TIB-152), HEK 293T (Cat#: CRL-3216), A549 (Cat#: CCL-185) and 293FT (Cat#: R70007) were obtained from ATCC. LP-1 (Cat#: ACC 41) cells were obtained from DSMZ-German Collection (Leibniz Institute, Germany). KMS-12 (Cat# JCRB0429) cells were from the Japanese Collection of Research Bioresources (National Institutes of Health Sciences, Japan). RPMI-8226 V10R cells were a gift from Robert Orlowski (MD Anderson). Human umbilical vein endothelial cells (HUVEC) were purchased from Lonza (Cat#: C2519A). RPMI-1640 and DMEM media were purchased from Fisher Scientific (Cat#: 11875119 and Cat#: 11885092). EGM-2 medium was purchased from Lonza (Cat#: CC-3162). All cell lines were independently authenticated using short tandem repeat profiling and tested for mycoplasma using PCR within 3 months of use. Clinical samples were obtained from multiple myeloma patients providing informed consent in accordance with the Declaration of Helsinki, as approved by the Johns Hopkins Medical Institutes Institutional Review Board. Mononuclear cells were isolated by density centrifugation (Ficoll–Paque; Pharmacia). Protein assay kit was purchased from Bio-Rad (Cat#: 5000006).

#### Antibodies

Anti-PARP-1 (Cat#: sc-7150), anti-GAPDH (Cat#: sc-47724), anti-β-actin (4967S), anti-P53 (Cat#: SC-126), anti-LDH (Cat#: SC-133123) were purchased from Santa Cruz Biotechnology; Anti-Caspase-3 (Cat#: 9662S), anti-Caspase-7 (Cat#: 12827), anti-Caspase 8 (Cat#: 9746S), anti-β-Catenin (Cat#: 8480S), anti-CHOP (Cat#: 2895S), anti-Lamin-B1 (Cat#: 12586S), anti- K48 linkage specific polyubiquitin (Cat#: 4289S) were purchased from Cell Signaling Technology; Anti-PREP (Cat#: AF4308-SP) was purchased from R&D Bio.

#### Cell Culture

All cells were grown at 37°C with 5% $CO_2$ in a humidified environment. HUVEC were grown in EGM-2 bullet kit media with 2% or 10% FBS and used between passage 2 and 8. NCI-H929, RPMI-8226, U266, LP-1, Pfeiffer, MM1.S, and Jurkat T cells were grown in RPMI-1640 media with the addition of 10% FBS. HEK 293T and A549 cells were grown in DMEM media with the addition of 10% FBS.

#### Resazurin Viability Assay

Cells were seeded in a 96-well plate (Costar) to the following density in 190 μL media: NCI-H929, RPMI-8226, U266, LP-1, Pfeiffer, MM1.S 20,000 cells/well, Jurkat T 10,000 cells/well; HEK 293T and A549 1,000 cells/well. To avoid edge effect, the outer circle of wells were filled with media only. Then drugs were added with proper dilutions to 0.1% DMSO in a total of 200 μL media (For adhesion cells, drugs were added after an overnight recovery). Following a 72 h incubation, cells were added with 10 μL/well of resazurin sodium salt solution in PBS to a final concentration of 10 μg/mL and the plates were incubated at 37 °C for 6 h before reading the fluorescence F (544nm Ex / 590nm Em) with a plate reader. Relative cell viability was calculated based on the formula: Relative viability

= ($F_{drug\_treated}$-$F_{media}$)/($F_{DMSO\_control}$–$F_{media}$). GraphPad Prism (v4.03) software was used to determine $IC_{50}$ values using a four-parameter logistic regression. Synergy analysis was performed using SynergyFinder 3.0.(*50, 51*)

**Cell Cycle Analysis**

NCI-H929 were seeded at $1 \times 10^6$ cells / 2mL in each well of a 6-well plate, and treated with drugs or vehicle control for 16 h. Cells were washed with PBS twice and pelleted at 500 × g. The pellet was resuspended in 0.5 mL PBS and added dropwise using a Pastuer pipet to 2 mL 75% ethanol in a 5 mL polystyrene tube being slowly agitated by a vortex. The cells were stored at 4 °C until staining. To do so, cells were pelleted at 500 × g, resuspended in 5 mL PBS, rested 60 seconds, pelleted again and washed in 5 mL PBS. The cell pellet was then resuspended in 0.5 mL staining solution (0.1% Triton-X-100, 0.2 mg/mL DNase free RNase A, and 0.02 mg/mL propidium iodide). Cells were allowed to stain for 30 min – 1 h prior for analysis. Propidium iodide incorporation was measured using a BD FACSCalibur™ Flow Cytometer. The percentage of cells in each cell cycle stage was determined with FlowJo.

**Western Blot Analysis**

For western blot analysis, NCI-H929 cells were seeded at $1 \times 10^6$ cells / 2mL in each well of a 6-well plate, and treated with drugs or vehicle control for designated time. Celle were then harvested and lysed with RIPA buffer plus protease inhibitor (cOmplete™ Protease Inhibitor Cocktail, Roche #04693116001). Cell lysates were subjected to SDS/ PAGE and then transferred to a nitrocellulose membrane. Membranes were first blocked in 5% (wt/vol) BSA in Tris-buffered saline plus 0.1% Tween 20 (TBST) at room temperature for 30 min and incubated with primary antibodies at 4 °C for overnight. Membranes were then washed three times with TBST and incubated with secondary antibodies at room temperature for another 1 h. Membranes were washed with TBST three times again and incubated with ECL substrate for 1.5 min at room temperature. Pictures were captured using a GeneSys Image Station.

**Cell Apoptosis Assay by Flow Cytometry**

NCI-H929 cells were plated at $5 \times 10^5$ cells/mL in a 12-well plate, treated with Rapaprotin for 16 hours. The cells were washed with cold PBS buffer, stained using Dead Cell Apoptosis Kit with Annexin V FITC and PI (Life Technology #V13242), and analyzed using BD FACSCelesta Flow Cytometer. Data was processed using Flowjo.

**Proteasome Inhibition Assay in Cell Lysates**

Assay was performed based on literature.(52) Lysates of NCI-H929 cells were prepared in proteasome lysis buffer (50 mM HEPES (pH 7.8), 10 mM NaCl, 1.5 mM $MgCl_2$, 1 mM EDTA, 1 mM EGTA, 250 mM sucrose, 5mM DTT) with sonication at $30 \times 10^6$ cells/mL and centrifuged 16,000 × g for 10 min at 4 °C to remove cell debris and supernatant was used fresh. Each 50 μL lysate was incubated in 200 μL proteasome assay buffer (lysis buffer with 2 mM ATP) for 60 min at 37 °C in a black walled 96-well plate with inhibitor at designated concentrations and 100 μM of fluorogenic substrate: Suc-LLVY-AMC (Enzo Life Sciences #P802) for chymotrypsin-like activity, Z-ARR-AMC (EMD Millipore #539149) for trypsin-like activity, or Z-LLE-AMC (EMD Millipore #539141) for caspaselike activity. Then, fluorescence signal F (360 nm Ex/ 460 nm Em) was read on fluorescent plate reader. Relative proteasome activity was calculated based on the formula: Relative proteasome activity = $(F_{Sample}-F_{media})/(F_{enzyme}-F_{media})$. GraphPad Prism (v4.03) software was used to determine $IC_{50}$ values using a four-parameter logistic regression.

**Proteasome Inhibition Assay with Recombinant 26S Proteasome**

Assay was modified based on literature.(53) Human 26S proteasome (10 nM, Boston Biochem # E-365) was incubated for 5 hours at 37 °C in a black-walled 384-well plate in 50 μL developing buffer (50 mM Tris, pH 7.5, 150 mM NaCl, 5 mM $MgCl_2$, 1 mM ATP, 1mM DTT, 0.1 mM ATPγS (Cayman Chemical #14957)) and 100 μM of fluorogenic substrate specified above. Then, fluorescence signal F (360 nm Ex/ 460 nm Em) was read on fluorescent plate reader. Relative proteasome activity was calculated based on the formula: Relative proteasome activity = $(F_{Sample}-F_{media})/(F_{enzyme}-F_{media})$.

**Generation of CRISPR/Cas9 knockout library in NCI-H929 cells**

The lenti-CRISPRv2 based human whole genome CRISPR Knockout Pooled Library (Brunello) Pooled Library (Addgene #73179) was amplified according to the protocol provided by Addgene(54) and yielded 129 copies per gRNA. The library was packaged into lentiviral particles using pMD2.G and psPax2 packaging vectors, 293FT feeder cells, and Fugene HD transfection reagent as previously described(55). Approximately 10 million log-phase NCI-H929 cells were transduced with virus with a MOI (multiplicity of infection) between 0.3 and 0.4 and selected in the presence of 0.25 μg/mL puromycin for 7 days to ensure all remaining cells are transduced with only one plasmid. After selection, puromycin was removed and cells were allowed to recover for 3 days.

**Genome-Wide CRISPR Screen for Rapaprotin Resistant Cells**

Cells transduced with Brunello library were treated with Rapaprotin for 5 days to achieve less than 0.01% viability, washed, and then plated in methylcellulose (without Rapaprotin) as previously described(55). After 10 days, visible colonies were isolated and expanded for characterization. To identify the sgRNA corresponding to each resistant colony, genomic DNA was isolated using QIAamp DNA Mini Kit (51304). gRNA containing fragment was amplified with PCR using PfuUltra II Fusion High-fidelity DNA polymerase (Agilent #600670) with forward primer 5'-GGGACAGCAGAGATCCAGTTTG-3' and reverse primer 5'- CCCACTCCTTTCAAGACCTAGCT-3'. PCR product was gel-extracted and sequenced with primer 5'- GTACAAAATACGTGACGTAGA-3'.

**CRISPR-Cas9 Knockout of Prolyl Endopeptidase (PREP)**

Guide RNAs targeting PREP were designed with CRISPR design tool (http://crispr.mit.edu) and sequences are listed below. gRNAs 2-4 were included in the Brunello library, while gRNA1 was not included in the library. gRNA1-4 sequences are shown in the following. gRNA1: CCCAACATACTGTCTGACGA; gRNA2: CATCGTCAGACAGTATGTTG; gRNA3: GAATGTTCTTGACGTCATGG; gRNA4: GCAGGAATCCAGTGGCATCG. gRNAs 1-4 were individually cloned into LentiviralV2 vector (Addgene #52961). PREP-KO NCI-H929 cells were generated following the above protocol. Efficiency of PREP knock out was evaluated by Western blot 10 days after transduction and puromycin selection using anti-PREP antibody.

**Generation of mDHFR-GFP Reporter Cell Line**

The destabilized dihydrofolate reductase mutant (mDHFR)-GFP gene was cloned into pLenti6M vector. The cloned gene was confirmed not to contain any spurious mutations by sequencing the full length of the cloned insert. The gene product was then transfected into HEK293T cells with pSPAX2 and pMD2G using lipofectamine 2000 and lentiviruses were harvested after 72 h. NCI-H929 cells were infected with the corresponding lentivirus and cells stably expressing mDHFR-GFP were selected with 10μg/ml blasticidin for two weeks and maintained at the same concentration of antibiotic for culture.

**Proteasome Reporter Assay**

mDHFR-GFP reporter cells (FZ120-H929) were plated at 50,000 cells/well in a flat-bottom 96-well plate, treated with 200 nM of Bortezomib or various concentrations of Rapaprotin for 8 hours. After treatment, cells were washed with and re-suspended in FACS buffer (PBS with 2mM EDTA, 0.5% BSA and 0.02% $NaN_3$). Population of GFP-negative and -positive cells were analyzed with flow cytometry using Cytek Aurora. Data was processed with Spectroflow. GraphPad Prism (v4.03) software was used to determine the $IC_{50}$ value using a four-parameter logistic regression.

**Measuring of Concentrations of Rapaprotin and Rapaprotin-L in cell cultures**

Cells were seeded in a 12-well plate to the following density in 1mL media: suspension cells 1.0 million cells/well, adhesion cells 0.33 million cells/well. Rapaprotin was added into the cell culture immediately (suspension cells) or after overnight culturing (adhesion cells) to a final concentration of 1 μM. Cells were incubated at 37 °C with 5% $CO_2$ for designated time periods. After incubation, media was removed (suspension cells were spun down to remove media), and cells were washed with PBS twice. Then cells were lysed in 50 μL of RIPA buffer. Rapaprotin and Rapaprotin-L were extracted from cell lysate or 50 μL of cell culturing media by adding 30 μL of saturated $ZnSO_4$ and 90 μL of acetonitrile, vortex for 30 seconds and centrifuge for 5 minutes. Acetonitrile phase was separated and analyzed using LC-MS (Agilent 6120 Single Quad) with a Selective Ion Monitoring (SIM) mode. HPLC was run with flow rate 1.0 mL/min, with a gradient of acetonitrile (with 0.1% formic acid) from 50-95% in water (with 0.1% formic acid). Mass intensity was calculated using the total peak area of ion extraction result for m/z 1334.6 (Rapaprotin +$H^+$) and m/z 1356.6 (Rapaprotin+$Na^+$) at 13.1 min to represent Rapaprotin, and total peak area of m/z 1352.6 (Rapaprotin-L+$H^+$) and m/z 1374.6 (Rapaprotin-L+$Na^+$) at 11.0 min to represent Rapaprotin-L. Concentrations of Rapaprotin and Rapaprotin-L in acetonitrile extracts (C1) from cell pellet were calculated based on standard curves generated using designated concentrations of Rapaprotin and synthetic Rapaprotin-L. Intracellular concentrations (C2) were estimated based on the formula: C2 = C1 × (50 μL / volume of cell pellet).

**Estimating solubility of rapafucins in RPMI 1640 with 10% FBS**

Rapafucins are diluted in RPMI 1640 with 10% FBS to a final concentration of 50uM. The mixture was incubated at 37°C for 24h, then vortexed and filtered through a 0.2 μm syringe filter. 100μL of filtered solution was extracted with 50uL of saturated ZnSO4 and 150uL of acetonitrile. The acetonitrile phase was separated and injected into HPLC-MS for quantitative analysis.

**Purification of the human 26S proteasome**

Human proteasomes were affinity purified on a large scale from a stable HEK293 cell line containing HTBH (hexa-histidine, TEV cleavage site, biotin, and hexahistidine)-tagged RPN11 (a gift from L. Huang, University of California, Irvine). The cells were Dounce-homogenized in a lysis buffer (50 mM PBS (pH 7.5), 10% glycerol, 5 mM $MgCl_2$, 0.5% NP-40, 5 mM ATP and 1 mM DTT) containing protease inhibitors. Lysates were cleared by centrifugation (20,000g, 30 min), then incubated with NeutrAvidin agarose resin (Thermo Scientific) for 3 h at 4 °C. The beads were washed with excess lysis buffer followed by wash buffer (50 mM Tris-HCl (pH 7.5), 1 mM $MgCl_2$ and 1 mM ATP). 26S proteasomes were cleaved from the beads using TEV protease (Invitrogen). The resin was removed by centrifugation and the supernatant was then further purified by gel filtration on a Superose 6 10/300 GL column at a flow rate of 0.15 ml/min in buffer (30 mM Hepes (pH 7.5), 60 mM NaCl, 1 mM $MgCl_2$, 10% glycerol, 0.5 mM DTT, 0.6 mM ATP). Gel-filtration fractions were concentrated to about 2 mg/ml and the buffer was exchanged to 50 mM Tris-HCl (pH 7.5), 100 mM NaCl, 1 mM ATP and 10% glycerol.

**Native PAGE Western blot for 26S and 20S detection in NCI-H929 cells**

NCI-H929 cells ($10^6$) were seeded and treated with rapaprotin (or 0.1% DMSO) at indicated time points (1,000,000 cells/mL). Cells were washed 3x with ice cold PBS and resuspended in 1 mL of PBS. Disuccinimidyl suberate (DSS) (Thermofisher Cat.# 21555) was added to a final concentration of 0.25 mM and incubated for 30 minutes at room temperature. 1 M Tris HCl (pH=7.5) quenching solution was added to each sample for a final concentration of 15 mM and incubated for a further 15 minutes at room temperature. Cells were washed 3x with PBS then resuspended in native lysis buffer (25 mM Tris (pH 7.4), 10 mM $MgCl_2$, 10% glycerol, 1 mM ATP, 1 mM DTT). Cells were then lysed via sonication (continuous, 15% amplitude, 15 seconds). Samples were clarified, treated with 5x native loading dye (250 mM Tris (pH7.4), 50% Glycerol, 0.007% xylene cyanol), then run on a 4% native PAGE gel (130 V, 120 minutes). Proteins were transferred to a PVDF membrane overnight (60 mA, 18 hours). Membranes were probed with anti-β5 subunit antibody (Enzo, Cat.# BML-PW8895-0025).

**Cryo-EM sample preparation and data collection.**

We prepared 1.5 mg/ml human proteasome solution by thawing sample stored at -80 °C. To remove the glycerol in the solution, the complex system was applied to Zeba Micro Spin Desalting Columns (7K, Thermo Fisher), exchanging the buffer to 50 mM Tris- HCl (pH 7.5) containing 100 mM NaCl, 2mM ATP and 5mM $MgCl_2$. The sample is then carried on ice to the cryo-plunging room where diluted LMC solution was added to achieve a LMC concentration 5 μM (DMSO final concentration: 0.05 %). The sample was incubated at room temperature for 15 minutes or 90 minutes (two different time-resolved experiments). In all cases, we added 0.005% NP-40 to the proteasome solution immediately before cryo-plunging.

Cryo-EM grids were prepared with FEI Vitrobot Mark IV. C-flat grids (R1/1 and R1.2/1.3, 300 Mesh, Protochips) were glow-discharged before a 2.5-μl drop of 1.5 mg/ml proteasome solution was applied to the grids in an environmentally controlled chamber with 100% humidity and temperature fixed at 4 °C. After 1 or 2 s of blotting, the grid was plunged into liquid ethane and then transferred to liquid nitrogen. CryoEM data was collected with good-quality grids in a 300 kV Titan Krios G2 microscope (Thermo Fisher) equipped with the post-column BioQuantum energy filter (Gatan) connected to a K2 Summit direct electron detector (Gatan). Coma-free alignment and parallel illumination were manually optimized prior to each data collection session. Cryo-EM data were acquired automatically using SerialEM software in a super-resolution counting mode with 20 eV energy slit, with the nominal defocus set in the range of −0.7 to −2.2 μm. A total exposure time of 10 s with 250 ms per frame resulted in a 40-frame movie per exposure with an accumulated dose of ~50 electrons per Å$^2$. For the 90-minute experiment, the calibrated physical pixel size and the super-resolution pixel size were 0.84 Å and 0.42 Å, respectively, and a total of 15,461 movies was collected. For the 15-minute experiment, the calibrated physical pixel size and the super-resolution pixel size were 1.37 Å and 0.685 Å, respectively, and a total of 14,488 movies was collected.

**Cryo-EM data processing**

For clarity, we use the 15-minute data set for the following description and list the differences for the 90-minute data set at the end. Drift correction and dose weighting were performed using the MotionCor2 program(56) at a super-resolution pixel size of 0.685 Å. Drift-corrected micrographs were used for the determination of the micrograph CTF parameters with the Gctf program(57). Particles were automatically picked on micrographs that were fourfold binned to a pixel size of 2.74 Å using an improved version of the DeepEM program(58). A total of 996,750 particles were picked. Reference-free 2D classification and 3D classification were carried out in software packages RELION(59) version 3.0 and ROME(60). Focused 3D classification, CTF and aberration refinement, and high-resolution auto-refinement were mainly done with RELION 3.0, whereas the AlphaCryo4D software was used to analyze the conformational changes.

Double capped proteasome particles were separated from singly capped ones through several rounds of 2D and 3D classification. These particles were aligned to the consensus models of the doubly and singly capped proteasome to obtain their approximate shift and angular parameters. With these parameters, each doubly capped particle was split into two pseudo-singly capped particles by re-centering the box onto the RP–CP subcomplex. Then the box sizes of pseudo-singly capped particles and true singly capped particles were both shrunk to 560 × 560 pixels with a pixel size of 0.685 Å, and down-sampled by two-fold to a pixel size of 1.37 Å for the following processing. A total of 1,594,832 particles were obtained after this step. Particles were aligned to the CP subcomplex through auto-refinement with a CP mask, followed by one round of CTF refinement.

In the following steps, the entire particles set are shuffled, classified, and voted using the AlphaCryo4D protocol(*30*). Specifically, the set was first divided into 8 equal parts and shuffled with parameter $M = 3$ to generated $8 \times (3+1) = 32$ particles groups. Each group is then classified into 30 classes, generating altogether 960 3D volumes. The features of these volume were learned using deep neural network (DNN) by the Adam algorithm of initial learning rate 0.01. Guided by these features represented in the DNN encoding layer, the volumes were embedded on a 2D surface (pseudo-energy landscape) using t-SNE algorithm. A few clusters of interest with high particle populations were voted to generate the particles sets that used in the subsequent auto-refine processes to generate higher resolution maps for the entire CP and RP complex.

To further classify the RP states, 938735 particles which contain intact RP were separated from the other particles through alignment-skipped 3D classification and AlphaCryo4D. These particles were then refined with the CP masked and the CP density was subtracted to eliminate the influence of CP on auto-refinement and classification. The particle box was recentered to the RP and shrunk to 240 × 240 pixels, with a pixel size of 1.37 Å. The CP-subtracted particles were aligned to the RP subcomplex. Alignment-skipped RP-masked 3D classification was then performed, followed by AlphaCryo4D analysis, which yielded different RP conformational states.

For the 90-minute data set, a similar data processing procedure was used except for the following differences. A total of 397,069 particles were picked from the raw micrographs, and 590,054 single cap pseudo particles entered AlphaCryo4D analysis, with a particle images pixel size of 1.68 Å and box size of 220 × 220. Particles were divided into 6 parts and shuffling was carried out using a parameter $M = 5$ to generate 36 particle groups and 432 volumes in total after 3D classifications.

**Data and materials availability:**
Cryo-EM density maps of the RPN1-ATPase-CP and ATPase-CP sub-assemblies and their corresponding atomic models are being deposited into the Electron Microscopy Data Bank (EMDB) (www.emdataresource.org) and the Protein Data Bank (PDB) (www.pdb.org).

**Chemical Synthesis**

**Synthetic Reagents**
Piperidine, N,N-diisopropylethylamine (DIPEA) were purchased from Alfa Aesar. Anhydrous pyridine was purchased from Acros. Solid support resin with 2-chlorotrityl chloride (Cat#: 03498) was purchased from Chem-Impex. HATU was purchased from ChemImpex. Fmoc protected amino acid building blocks were purchased from ChemImpex, Novabiochem or GL Biochem. Dichloromethane (DCM or $CH_2Cl_2$), methanol (MeOH), hexanes, ethyl acetate (EtOAc), 1,2-dichloroethane (DCE, anhydrous), N,N’- dimethylformamide (DMF, anhydrous), Hoveyda-Grubbs catalyst 2nd generation and all the other chemical reagents were purchased from Sigma-Aldrich.

**Instruments for Synthesis and Purification**
NMR spectra were recorded with Burker-400 and -500. High performance liquid chromatographic analyses were performed with Agilent LC-MS system (Agilent 1260 series, mass detector 6120 quadrupole). Orbital shaking for solid-phase reactions was performed on a Mettler-Toledo Bohdan MiniBlock system for 96 tubes (30-200mg resin in SiliCycle tubes) or a VWR Mini Shaker (0.2-2g resin in a plastic syringe with a fritted disc). Reagents were added with an adjustable Rainin 8-channel pipette for the MiniBlock system. Microwave reactions were performed with a Biotage Initiator Plus or Multiwave Pro with silicon carbide 24-well blocks from Anton Parr. Compound

purification at 0.05-50g scale was performed with Teledyne Isco CombiFlash Rf 200 or Biotage Isolera One systems followed by a Heidolph rotary evaporator followed by overnight drying with a custom-designed box (50 cm × 50 cm × 15 cm) that allows air flowing rapidly inside to remove the solvent. The high-throughput weighing of the compounds in the library was done by a Mettler-Toledo analytical balance that linked (Sartorious Entris line with RS232 port) to a computer with custom-coded electronic spreadsheet.

**Synthesis of Rapaprotin**

Rapaprotin was synthesized through solid-phase peptide synthesis (SPPS), followed by microwave-assisted ring-closing metathesis (RCM) reaction, deprotection of the N-methyl tyrosine and installation of the 4-morpholinecarbonyl moiety. Protocols for SPPS and RCM were adopted from reference (*1*). Fmoc-D-Phe-OH, Fmoc-Gly-OH, Fmoc-D-Phe-OH, Fmoc-N-Me-Tyr(tBu)-OH and eFKBD (FKBD11) were coupled sequentially on to C6-linker chlorotrityl resin (General Procedure A in reference (*1*)) before microwave-assisted RCM reaction (General Procedure B in reference (*1*)). Coupling reactions were monitored by cleaving compounds from the resin using 10% TFA in DCM, and analyzed with LC-MS. After RCM reaction, resin was removed with filtration, and TFA was added into reaction supernatant to 10% while stirring. The reaction was stirred at room temperature for 2 hours and monitored with LC-MS. When complete conversion was achieved, Rapaprotin B (off-white foaming solid, 10-30% final combined yield according to resin loading capacity) was obtained by silica gel purification with 0-30% solvent B (DCM/EtOAc/MeOH 2:2:1) in solvent A (DCM). HRMS $[M+H]^+$ found 1221.5760 (Calc. $[C_{68}H_{81}N_6O_{15}]^+$ = 1221.5760).

Rapaprotin B (1.01 g, 0.75 mmol) was dissolved in 25 mL dry DCM in a round bottom flask. Then pyridine (500 µL) and $Et_3N$ (230 µL, 2.0 eq.) was added via syringes. Then a solution of 4-Morpholinecarbonyl chloride (288 µL, 3.0 eq.) in 5mL dry DCM was added dropwise via a syringe. The reaction mixture was stirred under Ar at rt. for 2h. The progress was monitored by LC-MS. Then 50mL $NH_4Cl$ was added to the solution and stirred for 10min. 100mL EtOAc was added to extract the product. EtOAc phase was collected, and $H_2O$ phase was extracted once more with EtOAc. EtOAc phase was combined and dried with $Na_2SO_4$. Then solvent was evaporated and Rapaprotin (off-white foaming solid, 560 mg, 51%) was obtained by silica gel purification with 0-30% solvent B (DCM/EtOAc/MeOH 2:2:1) in solvent A (DCM) followed by reverse phase (C18) flash column purification with 0-95% acetonitrile in $H_2O$. HRMS $[M+H]^+$ found 1334.6239 (Calc. $[C_{73}H_{88}N_7O_{17}]^+$ = 1334.6237).

**Synthesis of Rapaprotin-L**

*Chlortrityl Resin (S,E)-1-(4-((5-((((9H-fluoren-9-yl)methoxy)carbonyl)(methyl)amino)pent-2-enoyl)oxy)-3,3-dimethyl-2-oxobutanoyl)piperidine-2-carboxylate* **(1)** 115 mg of Cl-Tr-Resin (Chem-Impex, 0.19 mmol) was shaken with DCM (4 ml) in a 5 ml reaction vessel for 30 minutes. Filtered to remove solvent. Shaken with DMF for 30 minutes. Filtered to remove solvent. Resin was treated with (S,E)-1-(4-((5-((((9H-fluoren-9-yl)methoxy)carbonyl)(methyl)amino)pent-2-enoyl)oxy)-3,3-dimethyl-2-oxobutanoyl)piperidine-2-carboxylic acid (56.0 mg, 1 Eq,

94.8 µmol) in DMF (2 mL) and DIEA (66.2 mg, 89 µL, 5.4 Eq, 512 µmol) in DMF (2 mL) , shake for 3 hours. Filtered to remove reagents. Resin was treated with methanol (4 ml) and shaken for 30 minutes. Resin was filtered to remove solvent, washed with DCM (5×5 ml) and dried under vacuum to give 231 mg of yellow resin (LC = 0.82 mmol/g).

*Chlortrityl Resin (2S)-1-((5S,8R,14R)-5,8,14-tribenzyl-1-(9H-fluoren-9-yl)-4,10,16,24,24-pentamethyl-3,6,9,12,15,21,25-heptaoxo-19-(piperidin-1-yl)-2,22-dioxa-4,7,10,13,16-pentaazahexacosan-26-oyl)piperidine-2-carboxylate (6) and Chlortrityl Resin methyl (S)-1-((5S,8R,14R,E)-5,8,14-tribenzyl-1-(9H-fluoren-9-yl)-4,10,16,24,24-pentamethyl-3,6,9,12,15,21,25-heptaoxo-2,22-dioxa-4,7,10,13,16-pentaazahexacos-19-en-26-oyl)piperidine-2-carboxylate* **(6a)** The title compounds were synthesized from 231 mg of Chlortrityl Resin (S,E)-1-(4-((5-((((9H-fluoren-9-yl)methoxy)carbonyl)(methyl)amino)pent-2-enoyl)oxy)-3,3-dimethyl-2-oxobutanoyl)piperidine-2-carboxylate (0.19 mmol) using standard Fmoc Solid Phase Peptide Synthesis (SPPS) to give a yellow resin. A small portion of resin was cleaved with 20 % TFA (DCM) to verify synthesis. MW $[M+H]^+$ 1117.5/1202.5 (Calc. $[C_{64}H_{73}N_6O_{12}]^+ = 1117.5$; $[C_{69}H_{84}N_7O_{12}]^+ = 1202.5$).

*Chlortrityl Resin (S)-1-((4S,7R,13R,E)-4,7,13-tribenzyl-1-(3-(3-(3,4-dimethoxyphenyl)propanoyl)phenoxy)-3,9,15,23,23-pentamethyl-2,5,8,11,14,20,24-heptaoxo-21-oxa-3,6,9,12,15-pentaazapentacos-18-en-25-oyl)piperidine-2-carboxylate (8) and Chlortrityl Resin (2S)-1-((4S,7R,13R)-4,7,13-tribenzyl-1-(3-(3-(3,4-dimethoxyphenyl)propanoyl)phenoxy)-3,9,15,23,23-pentamethyl-2,5,8,11,14,20,24-heptaoxo-18-(piperidin-1-yl)-21-oxa-3,6,9,12,15-pentaazapentacosan-25-oyl)piperidine-2-carboxylate* (**8a**) The title compounds were synthesized from a mixture of **6** and **6a** and 2-(3-(3-(3,4-dimethoxyphenyl)propanoyl)phenoxy)acetic acid (**7**) using SPPS. A small portion of resin was cleaved with 20 % TFA (DCM) to verify synthesis. MW $[M+H]^+$ 1221.5/1306.6 (Calc. $[C_{68}H_{81}N_6O_{15}]^+ = 1221.4$; $[C_{73}H_{92}N_7O_{15}]^+ = 1202.5$).

*(S)-1-((4S,7R,13R,E)-4,7,13-tribenzyl-1-(3-(3-(3,4-dimethoxyphenyl)propanoyl)phenoxy)-3,9,15,23,23-pentamethyl-2,5,8,11,14,20,24-heptaoxo-21-oxa-3,6,9,12,15-pentaazapentacos-18-en-25-oyl)piperidine-2-carboxylic acid* (**8+**)**:** The resin linked peptide (8 and 8a, 0.0433 mmol) was swelled with 2 ml of DCM for 15 minutes. A solution of mCPBA (113 mg, 0.655 mmol) in 2 ml of DCM was added. shake for 30 minutes. Drained and washed with 3x2ml DCM. Treated with TFA (0.4 ml) in 2 ml of DCM for 2 minutes. Washed with MeOH and filtered. Mother liquor was reduced under vacuum to give a brown solid (38.4 mg, 72.6 %). MW $[M+H]^+$ 1221.5 (Calc. $[C_{68}H_{81}N_6O_{15}]^+ = 1222.4$).

*(S)-1-((4S,7R,13R,E)-4,7,13-tribenzyl-1-(3-((R)-3-(3,4-dimethoxyphenyl)-1-hydroxypropyl)phenoxy)-3,9,15,23,23-pentamethyl-2,5,8,11,14,20,24-heptaoxo-21-oxa-3,6,9,12,15-pentaazapentacos-18-en-25-oyl)piperidine-2-carboxylic acid* (**Rapaprotin-L**) Dichloro(pentamethylcyclopentadienyl)rhodium(III) dimer {[Cp*RhCl2]2} (1.94 mg, 0.10 Eq, 3.14 µmol) and (1R,2R)-(-)-N-p-Tosyl-1,2-diphenylethylenediamine {(R,R)-TsDPEN} (2.53 mg, 0.22 Eq, 6.92 µmol) were dissolved in water (1ml) while heating to 40 °C. (S)-1-((4S,7R,13R,E)-4,7,13-tribenzyl-1-(3-(3-(3,4-

dimethoxyphenyl)propanoyl)phenoxy)-3,9,15,23,23-pentamethyl-2,5,8,11,14,20,24-heptaoxo-21-oxa-3,6,9,12,15-pentaazapentacos-18-en-25-oyl)piperidine-2-carboxylic acid (38.4 mg, 1 Eq, 31.4 µmol) was dissolved in THF (2 ml), water (~2 ml) and sodium formate (21.4 mg, 10 Eq, 314 µmol) were added, let stir for 5 minutes. Catalyst solution was added, and mixture was heated to 40 °C for 3 hours, an additional 10 mg of sodium formate (0.157 mmol) was added, let Stir for 2 hours. Diluted with EtOAc. Layers separated. Washed with water (2x) and brine. Dried ($MgSO_4$) and reduced. Purified via PrepHPLC (5-95 % ACN/H2O) to give a white solid (21.2 mg, 55.0 %). MW $[M+H]^+$ 1352.6 (Calc. $[C_{73}H_{90}N_7O_{18}]^+$ = 1352.6). HRMS $[M+H]^+$ found 1352.6351 (Calc. $[C_{73}H_{90}N_7O_{18}]^+$ = 1352.6342).

Rapaprotin-B HRMS

**Elemental Composition Report** Page 1

**Single Mass Analysis**
Tolerance = 5.0 PPM / DBE: min = -1.5, max = 100.0
Element prediction: Off
Number of isotope peaks used for i-FIT = 3

Monoisotopic Mass, Even Electron Ions
589 formula(e) evaluated with 3 results within limits (up to 50 closest results for each mass)
Elements Used:
C: 0-200 H: 0-200 N: 5-9 O: 14-18 Na: 0-1
E269-72
Qtof_63180 68 (2.876) AM (Top,4, Ar,14000.0,716.46,0.70,LS 3); Cm (68:69) 1: TOF MS ES+ 1.33e+003

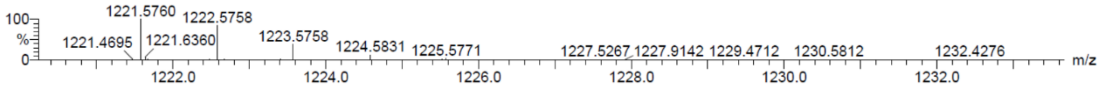

| | | | | | | |
|---|---|---|---|---|---|---|
| Minimum: | | | | -1.5 | | |
| Maximum: | | 5.0 | 5.0 | 100.0 | | |
| Mass | Calc. Mass | mDa | PPM | DBE | i-FIT | Formula |
| 1221.5760 | 1221.5760 | 0.0 | 0.0 | 31.5 | 2.1 | C68 H81 N6 O15 |
| | 1221.5736 | 2.4 | 2.0 | 28.5 | 4.7 | C66 H82 N6 O15 Na |
| | 1221.5720 | 4.0 | 3.3 | 27.5 | 8.4 | C63 H81 N8 O17 |

Rapaprotin-B HPLC

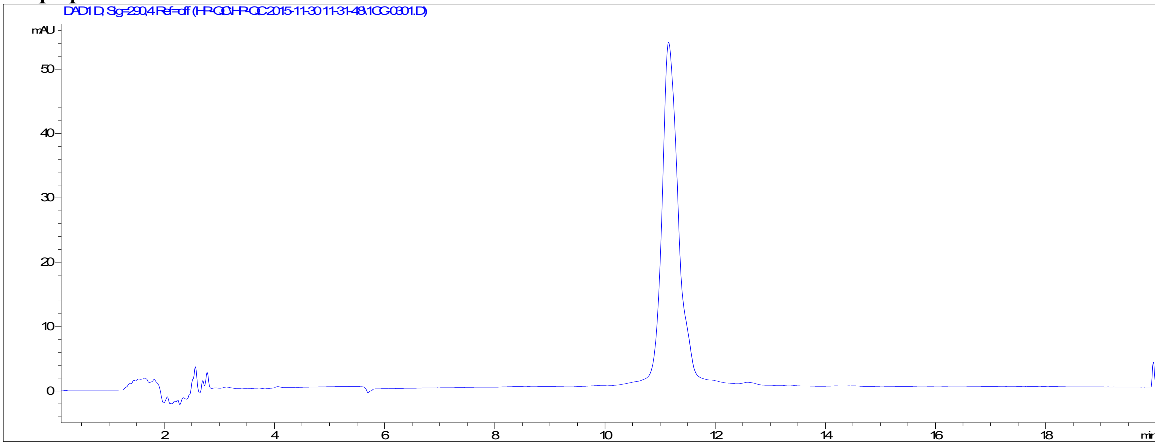

Rapaprotin-B [1]HNMR

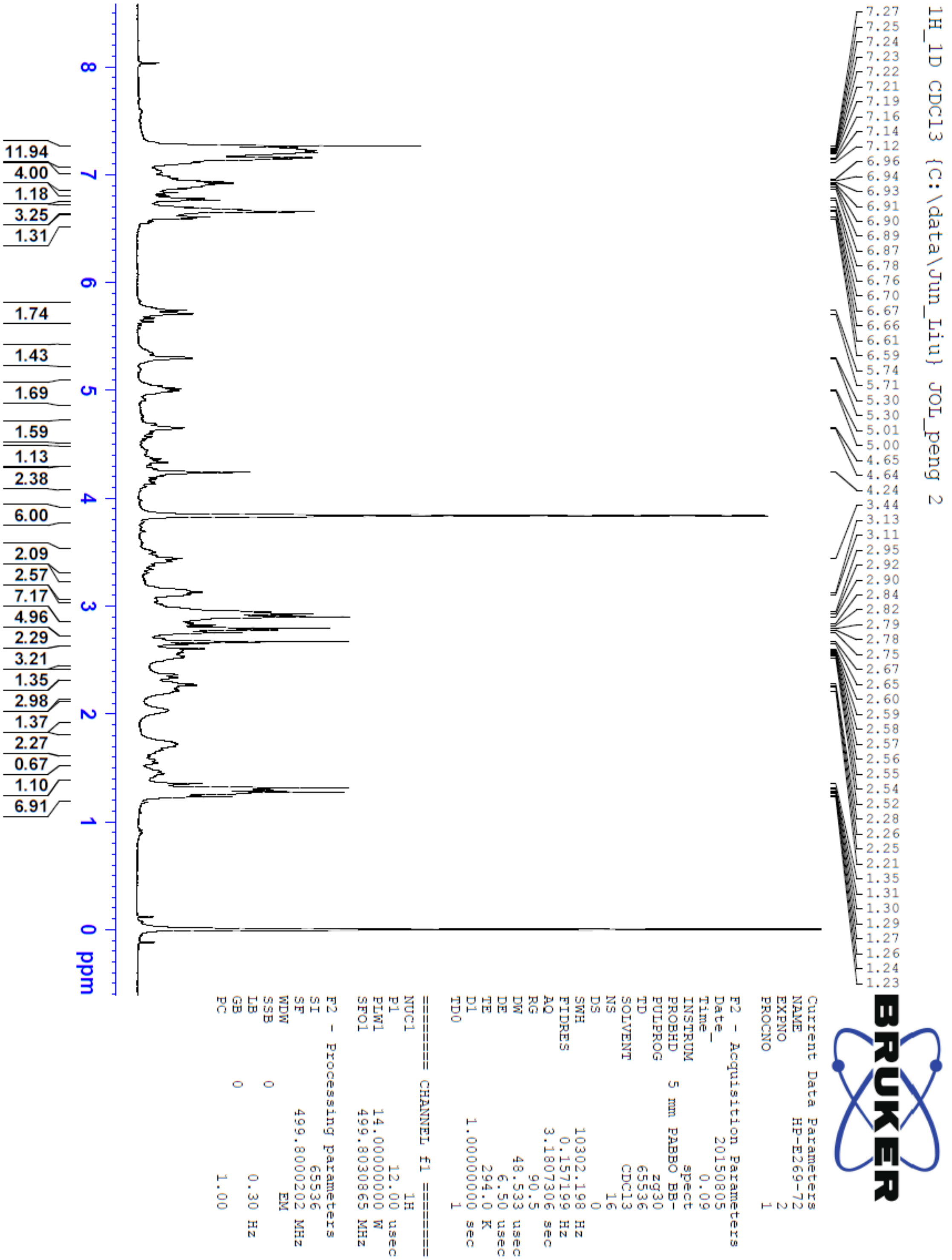

Rapaprotin-B $^{13}$CNMR, APT experiment

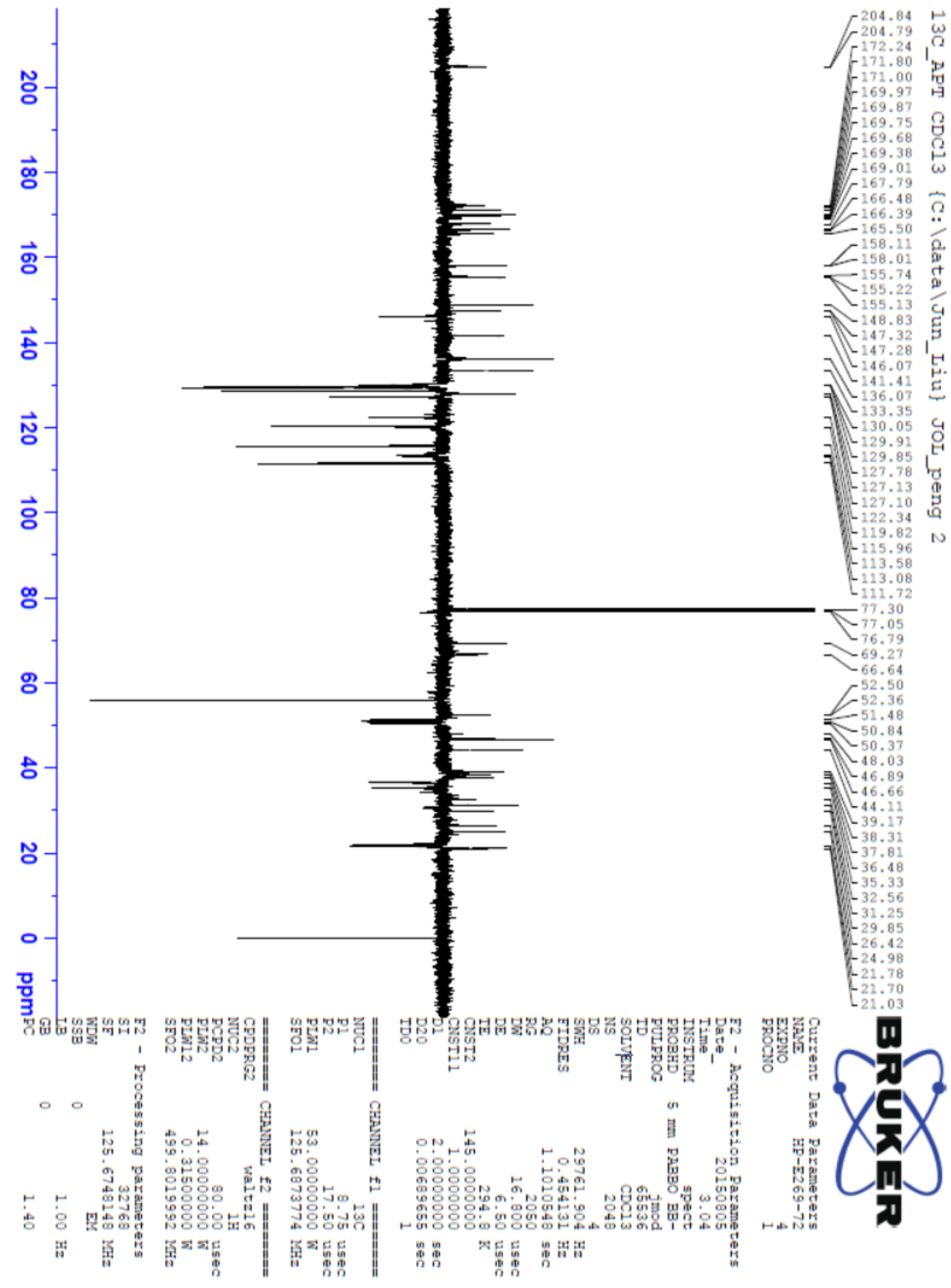

Rapaprotin HRMS

**Elemental Composition Report** **Page 1**

**Single Mass Analysis**
Tolerance = 5.0 PPM / DBE: min = -1.5, max = 200.0
Element prediction: Off
Number of isotope peaks used for i-FIT = 9

Monoisotopic Mass, Even Electron Ions
271 formula(e) evaluated with 4 results within limits (up to 50 closest results for each mass)
Elements Used:
C: 0-80 H: 0-100 N: 0-8 O: 8-19
Hanjing Peng E26972-MP MSL, School of Chemical Sciences, UIUC
Synapt2_9615a 19 (0.397) Cm (18:20-5:8) 1: TOF MS ES+ 2.23e+004

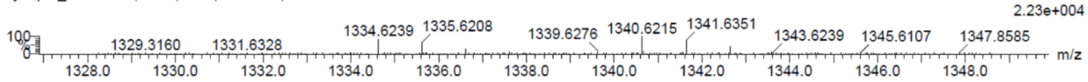

| | | | | |
|---|---|---|---|---|
| Minimum: | | | | -1.5 |
| Maximum: | | 5.0 | 5.0 | 200.0 |

| Mass | Calc. Mass | mDa | PPM | DBE | i-FIT | Norm | Conf(%) | Formula |
|---|---|---|---|---|---|---|---|---|
| 1334.6239 | 1334.6237 | 0.2 | 0.1 | 33.5 | 1120.1 | 1.176 | 30.86 | C73 H88 N7 O17 |
| | 1334.6264 | -2.5 | -1.9 | 32.5 | 1120.7 | 1.760 | 17.20 | C77 H92 N O19 |
| | 1334.6277 | -3.8 | -2.8 | 37.5 | 1120.7 | 1.808 | 16.40 | C78 H88 N5 O15 |
| | 1334.6178 | 6.1 | 4.6 | 42.5 | 1120.0 | 1.035 | 35.53 | C80 H84 N7 O12 |

Rapaprotin HPLC

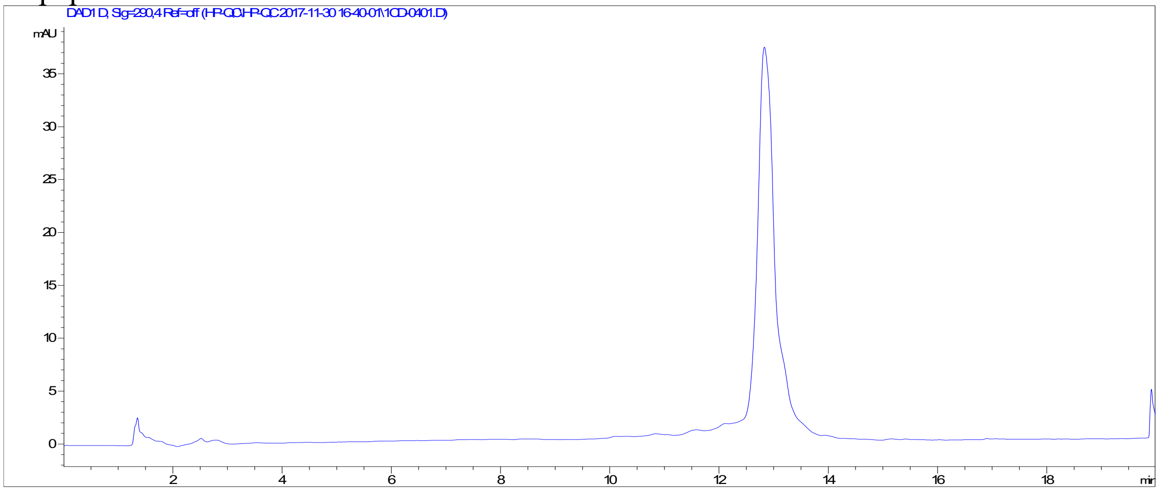

Rapaprotin $^{1}$HNMR

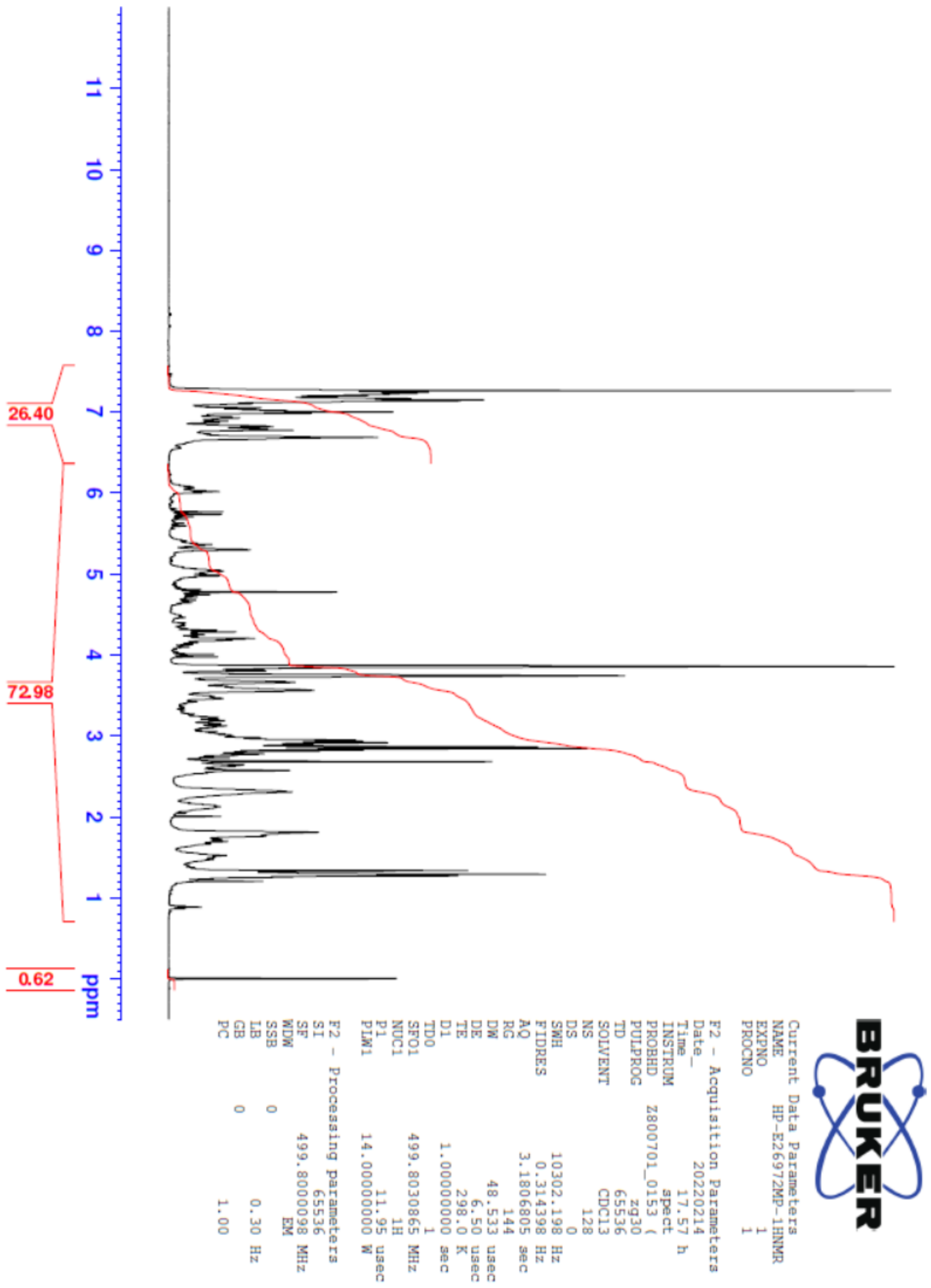

Rapaprotin $^{13}$CNMR, APT experiment

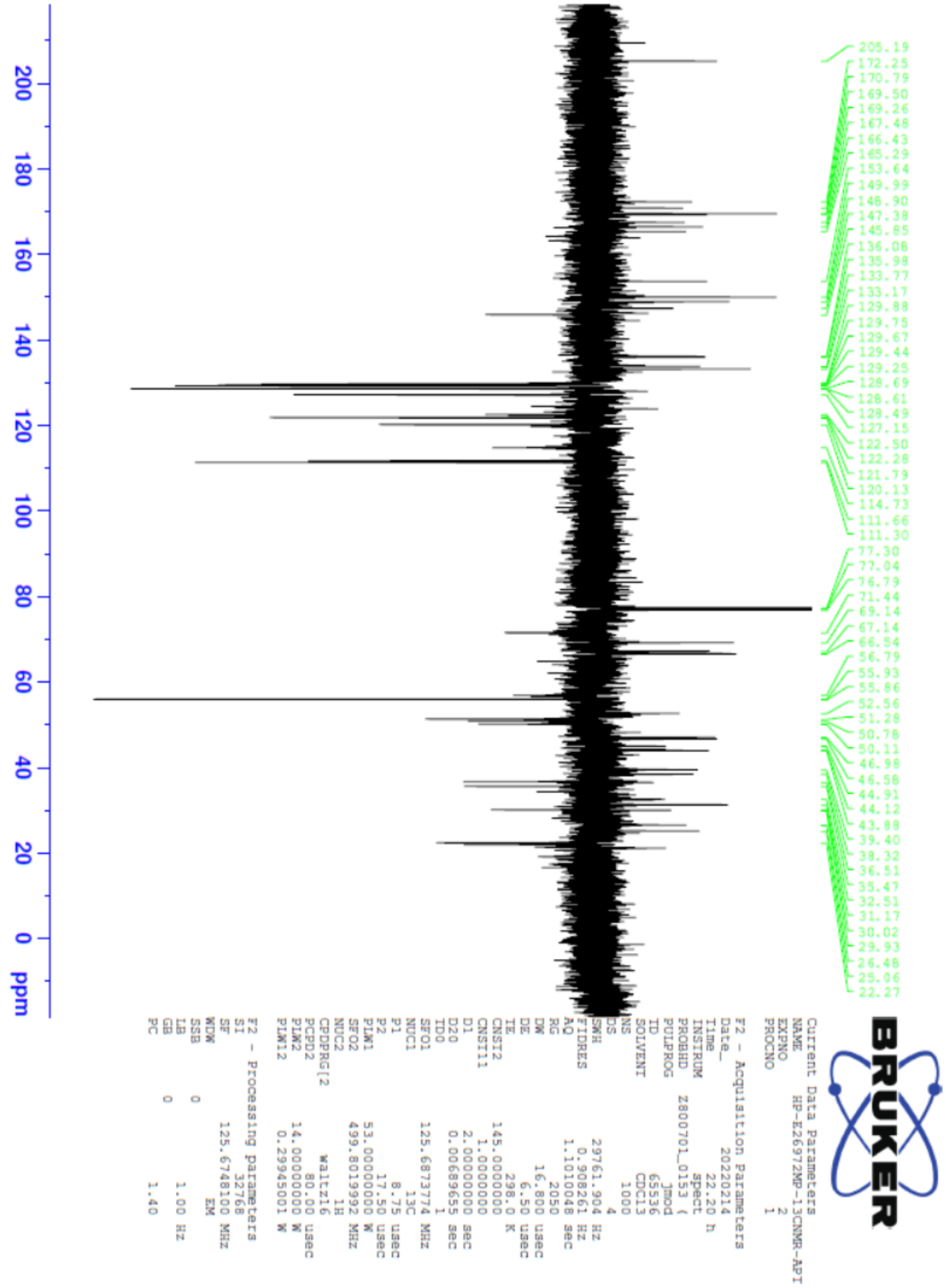

Rapaprotin NMR HSQC experiment

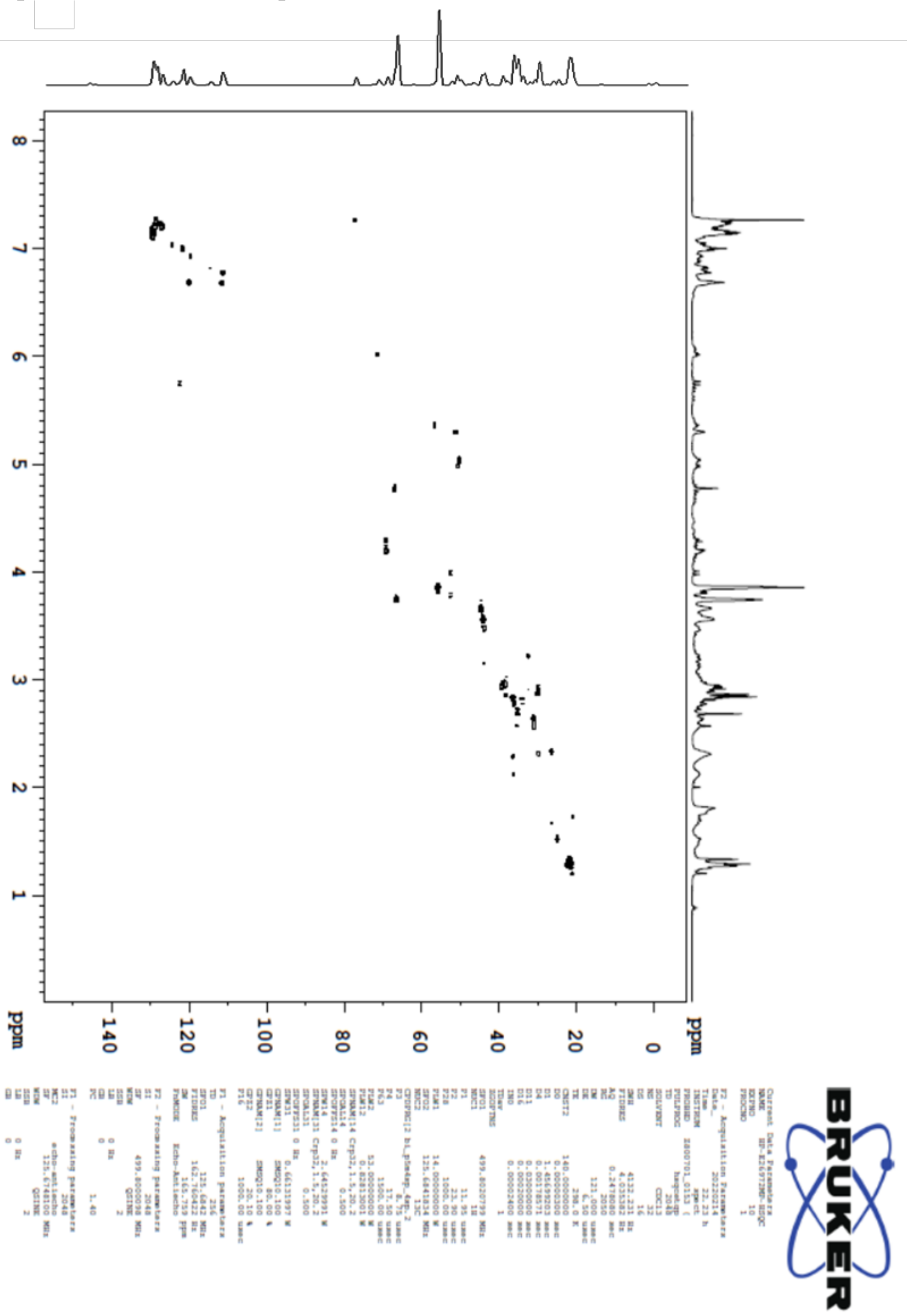

Rapaprotin-L HRMS

**Elemental Composition Report** **Page 1**

**Single Mass Analysis**
Tolerance = 5.0 PPM / DBE: min = -1.5, max = 200.0
Element prediction: Off
Number of isotope peaks used for i-FIT = 9

Monoisotopic Mass, Even Electron Ions
211 formula(e) evaluated with 3 results within limits (up to 50 closest results for each mass)
Elements Used:
C: 0-80 H: 0-100 N: 0-8 O: 8-19
Hanjing Peng Bru-018-018 MSL, School of Chemical Sciences, UIUC
Synapt2_9612c 18 (0.380) Cm (17:20-4:8) 1: TOF MS ES+
7.44e+004

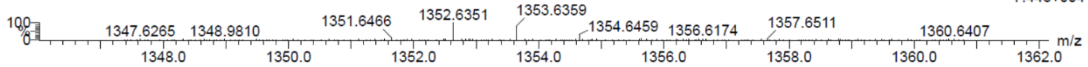

| Mass | Calc. Mass | mDa | PPM | DBE | i-FIT | Norm | Conf(%) | Formula |
|---|---|---|---|---|---|---|---|---|
| Minimum: | | | | -1.5 | | | | |
| Maximum: | | 5.0 | 5.0 | 200.0 | | | | |
| 1352.6351 | 1352.6342 | 0.9 | 0.7 | 32.5 | 1259.4 | 0.550 | 57.68 | C73 H90 N7 O18 |
| | 1352.6383 | -3.2 | -2.4 | 36.5 | 1260.3 | 1.430 | 23.94 | C78 H90 N5 O16 |
| | 1352.6284 | 6.7 | 5.0 | 41.5 | 1260.5 | 1.694 | 18.38 | C80 H86 N7 O13 |

Rapaprotin-L HPLC

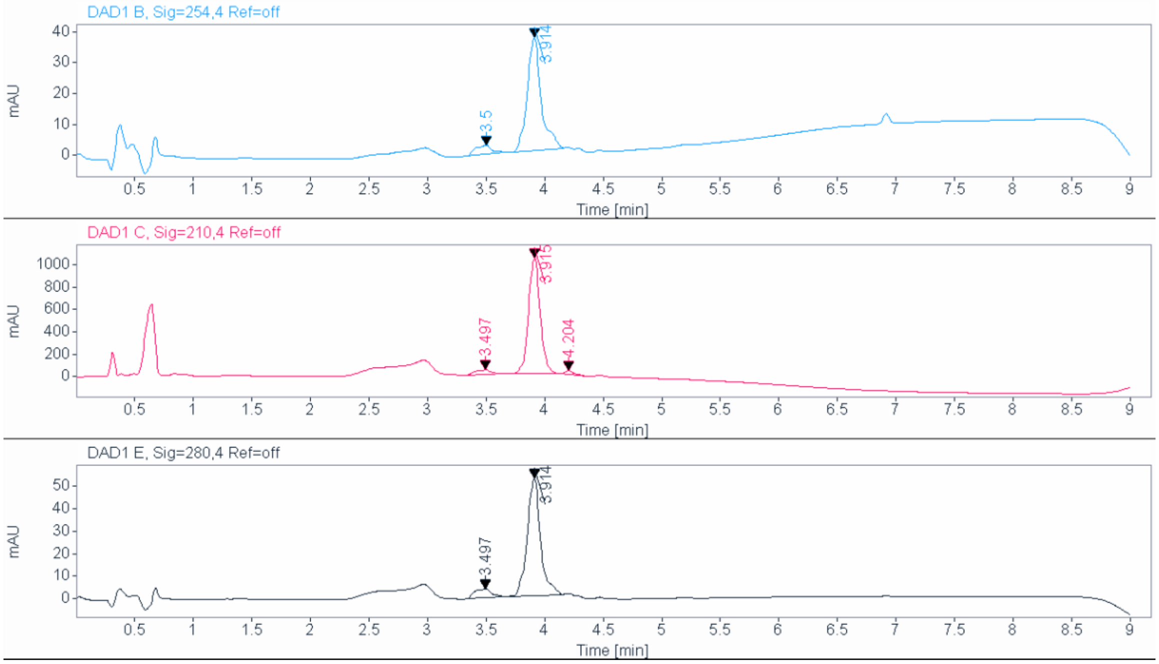

**Uncropped/full-size gels and blots:**

Uncropped blot for Figure 1f and 1g:

| | Left 7 lanes (excluding marker) | Right 6 lanes (excluding marker) |
|---|---|---|
| Blot | PARP and cleaved PARP for 1f | PARP and cleaved PARP for 1g |

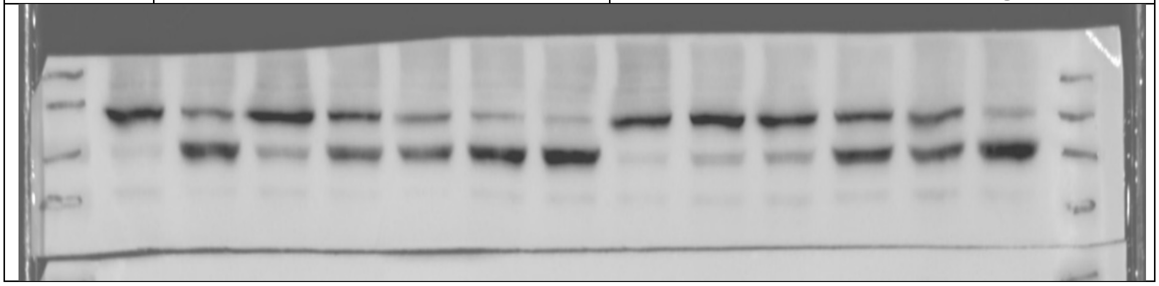

| | Left 7 lanes (excluding marker) | Right 6 lanes (excluding marker) |
|---|---|---|
| Blot | Cleaved PARP for 1f | Cleaved PARP for 1g |

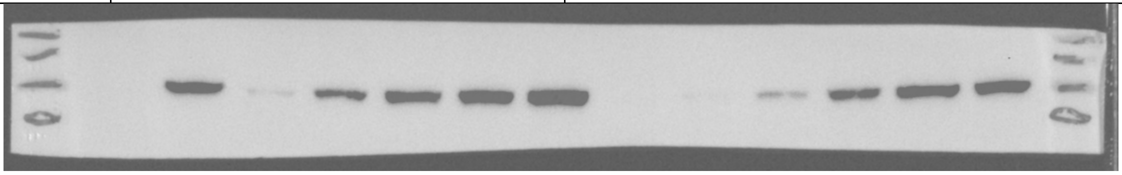

| | Left 7 lanes (excluding marker) | Right 6 lanes (excluding marker) |
|---|---|---|
| Blot | Caspase 3 and cleaved Caspase 3 for 1f | Caspase 3 and cleaved Caspase 3 for 1g |

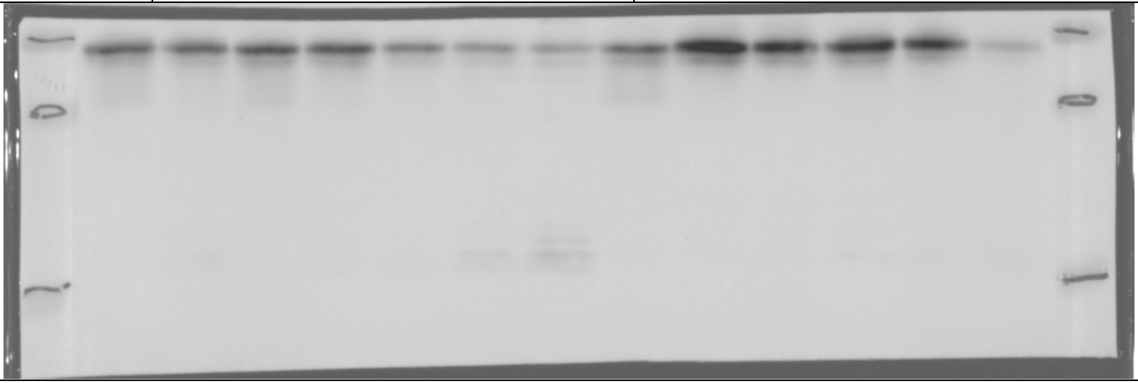

| | Left 7 lanes (excluding marker) | Right 6 lanes (excluding marker) |
|---|---|---|
| Blot | GAPDH for 1f | GAPDH for 1g |

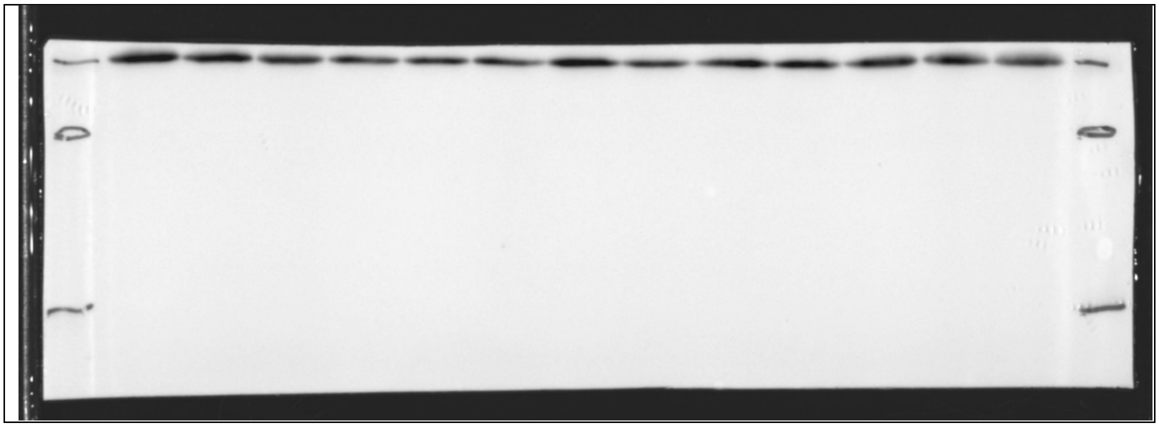

| | Left 7 lanes (excluding marker) | Right 6 lanes (excluding marker) |
|---|---|---|
| Blot | Caspase 7 and cleaved Caspase 7 for 1f | Caspase 7 and cleaved Caspase 7 for 1g |

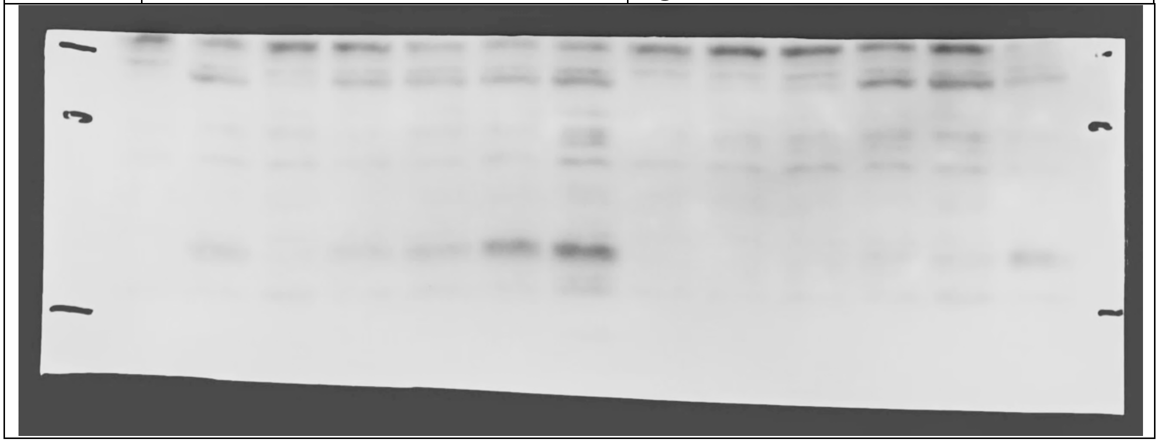

| | Left 7 lanes (excluding marker) | Right 6 lanes (excluding marker) |
|---|---|---|
| Blot | GAPDH for 1f | GAPDH for 1g |

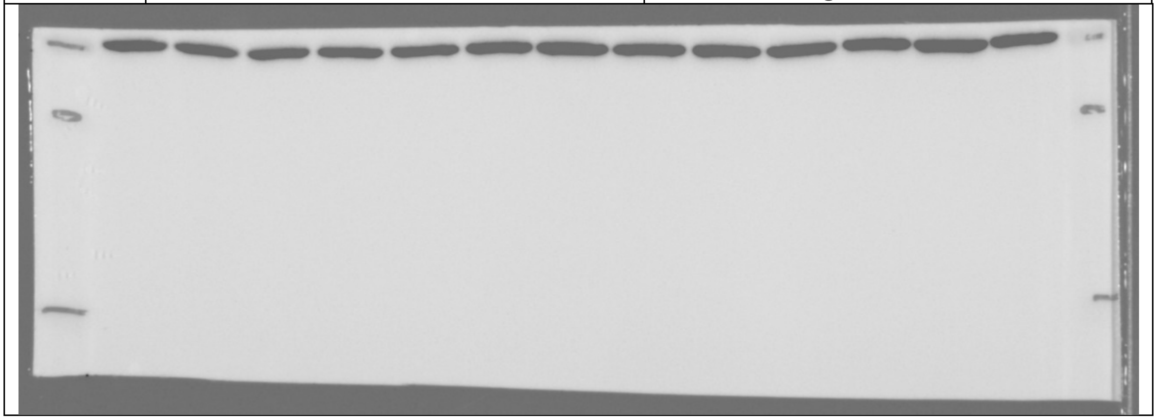

Uncropped blot for Figure 1h:

| Blot | Caspase 3 and cleaved Caspase 3 |
|---|---|

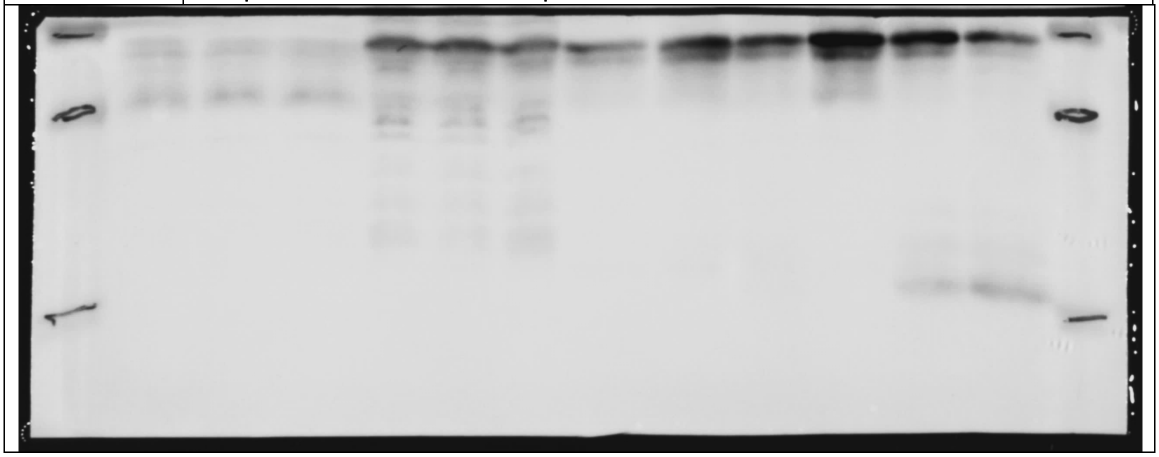

| Blot | HSP90 |
|---|---|

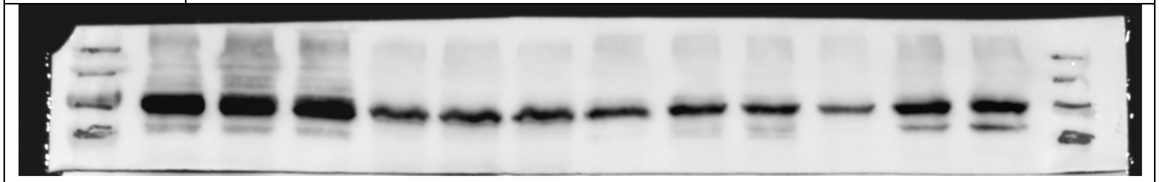

| Blot | GAPDH |
|---|---|

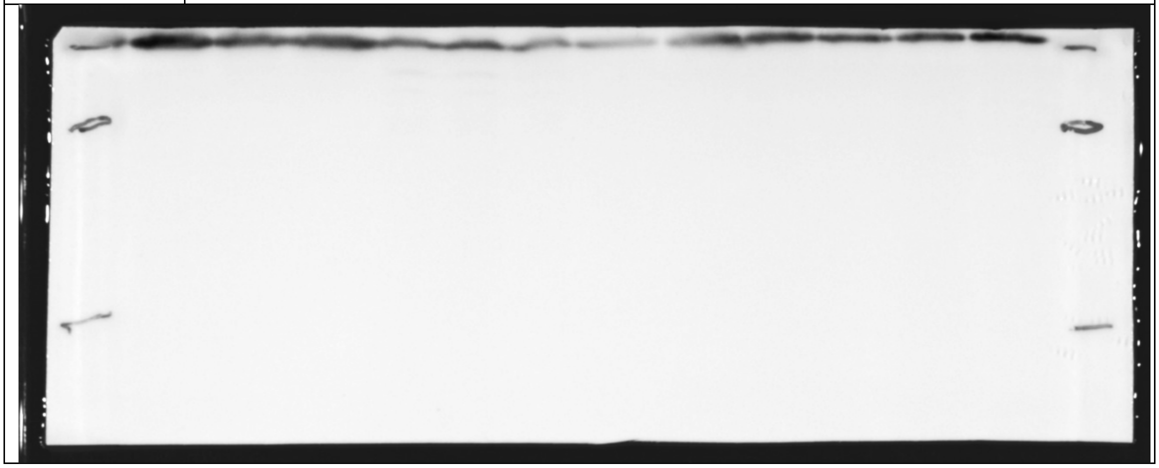

Uncropped blot for Figure 2c and 2f:

| | Left 6 lanes (excluding marker) | Lanes 2 and 3 from the right |
|---|---|---|
| Blot | PREP for 2f | PREP for 2c |

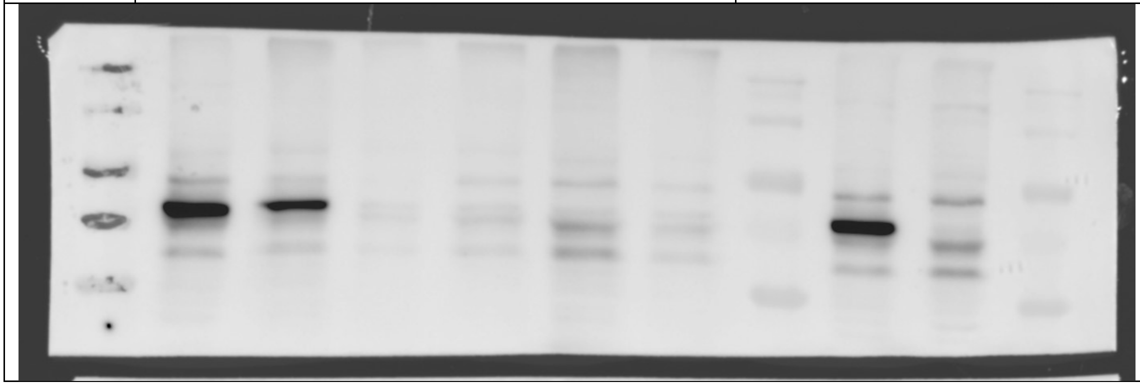

| | Left 6 lanes (excluding marker) | Lanes 2 and 3 from the right |
|---|---|---|
| Blot | GAPDH for 2f | GAPDH for 2c |

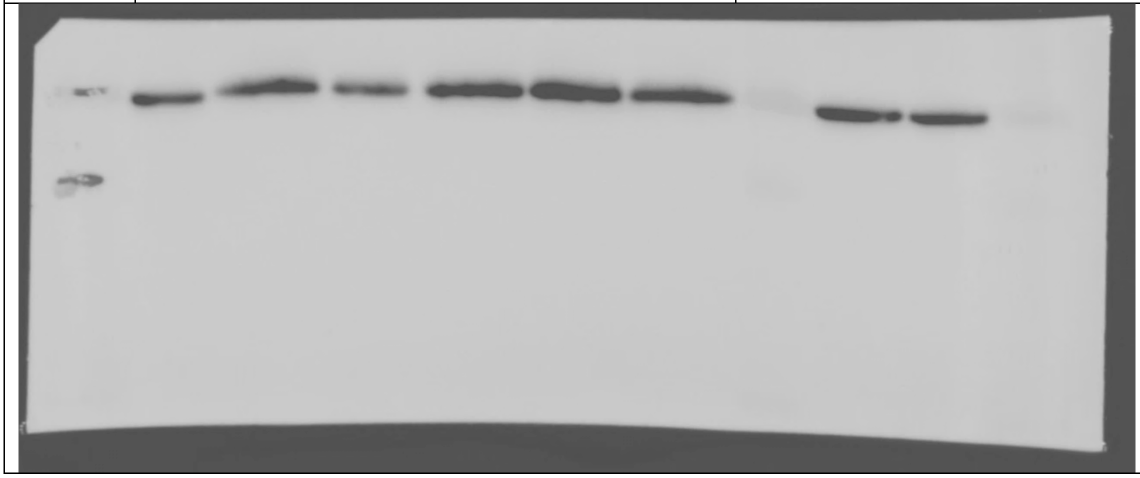

Uncropped blot for Figure 2d:

| | Left 6 lanes (excluding marker) |
|---|---|
| Blot | PARP and Cleaved PARP |

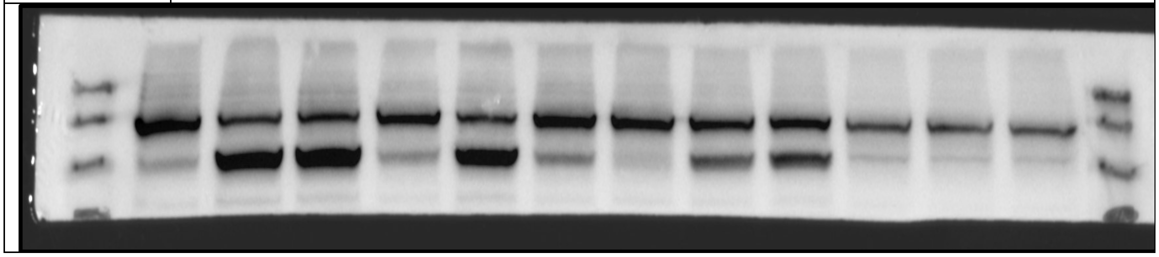

| | Left 6 lanes (excluding marker) |
|---|---|
| Blot | Caspase 3 and Cleaved Caspase 3 |

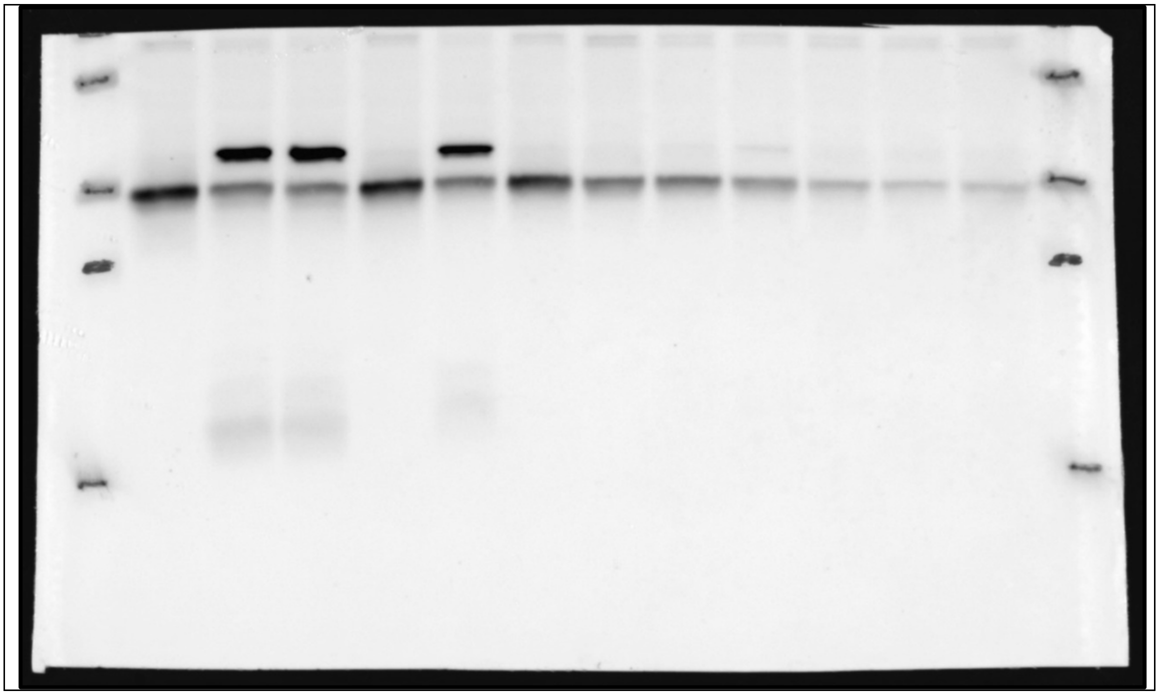

| | Left 6 lanes (excluding marker) |
|---|---|
| Blot | GAPDH |

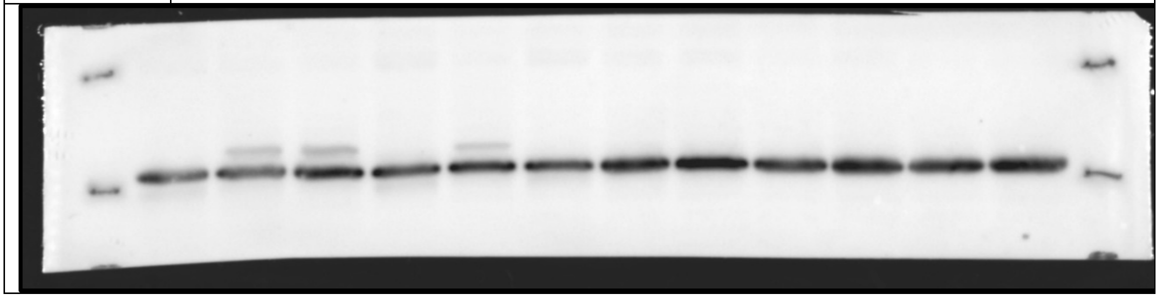

Uncropped blot for Figure 4d:

| Blot | K48-linked Ub |
|---|---|

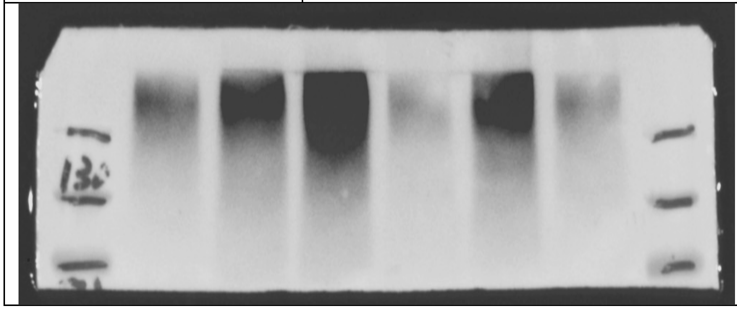

| Blot | GAPDH |
|---|---|

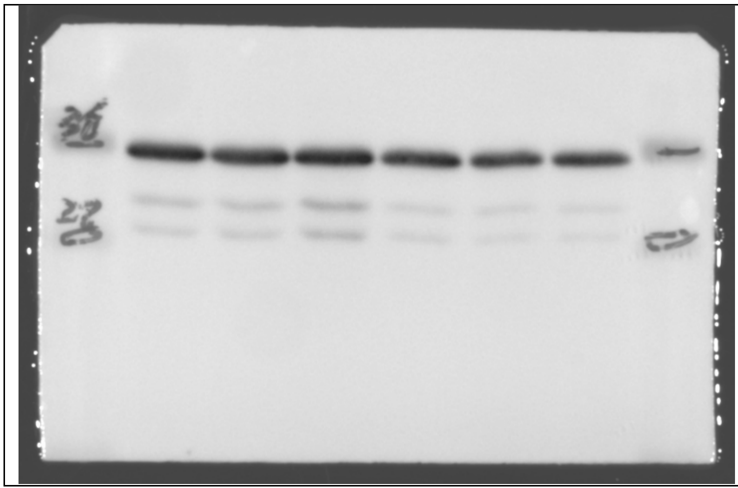

Uncropped blot for Figure 4e:

| Blot | CHOP |
|---|---|

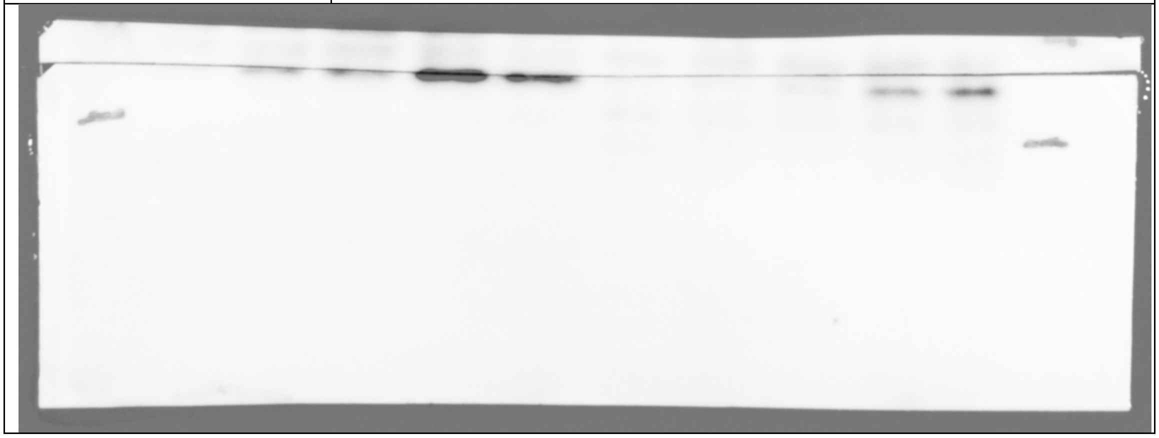

| Blot | Lamin-B1 |
|---|---|

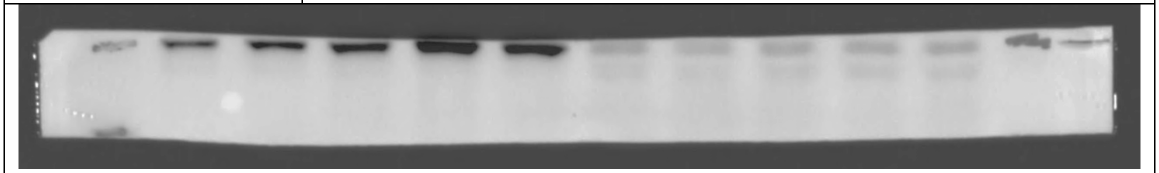

| Blot | LDH |
|---|---|

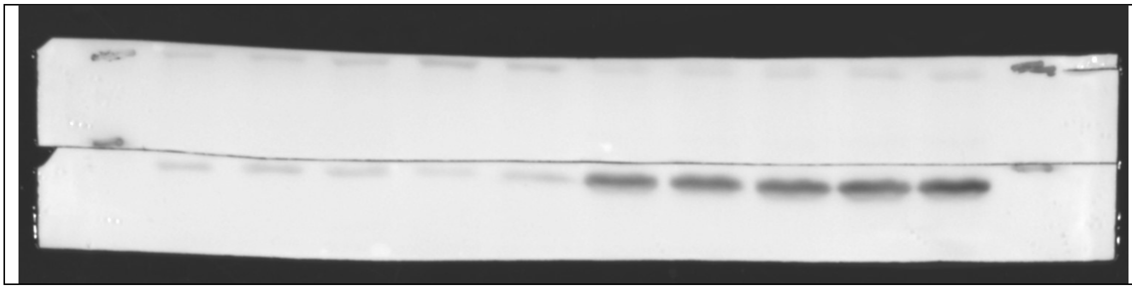

Uncropped gel for Figure 5d:

| Native gel probed for β5 subunit | Left 3 lanes |
| --- | --- |

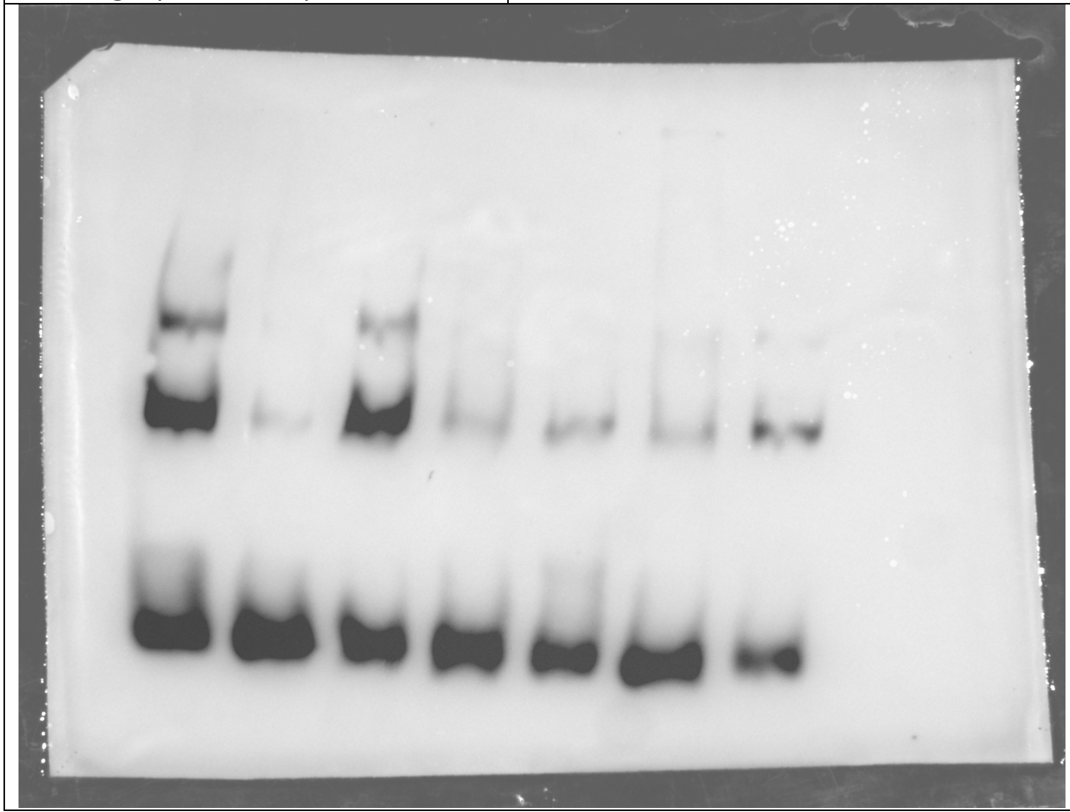

Uncropped blot for Figure S1f:

| | Left 7 lanes (excluding marker) |
| --- | --- |
| Blot | Caspase-8 and Cleaved Caspase-8 |

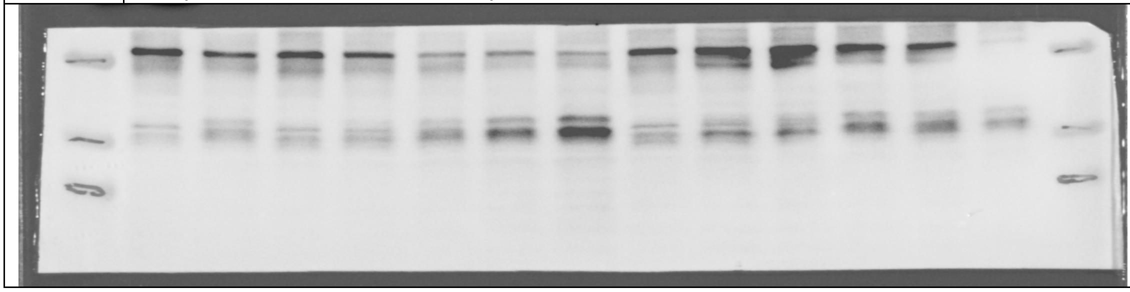

| | Left 7 lanes (excluding marker) |
|---|---|
| Blot | GAPDH |

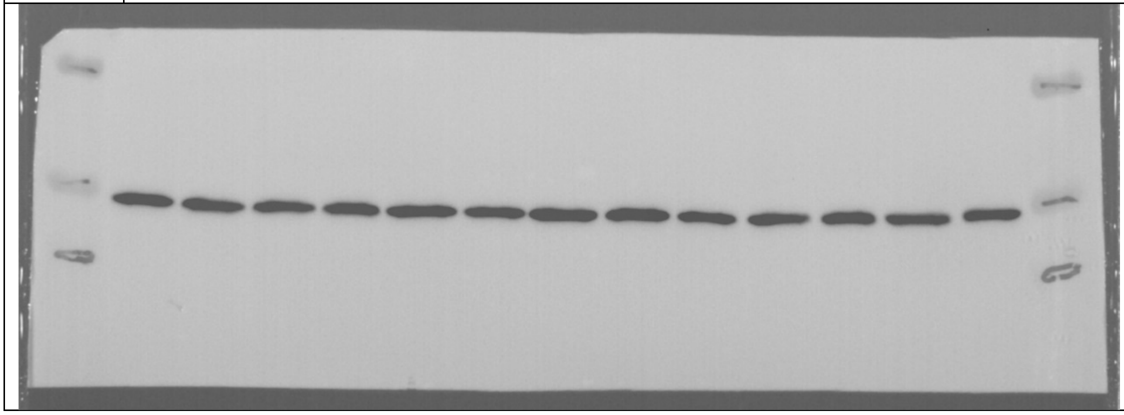

Uncropped blot for Figure S4a and S4b:

| | Left 6 lanes (excluding marker) | Right 6 lanes (excluding marker) |
|---|---|---|
| Blot | GAPDH for S4a | GAPDH for S4b |

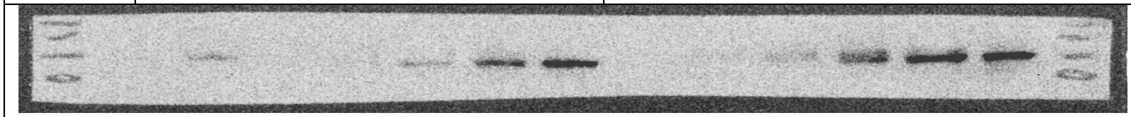

| | Left 6 lanes (excluding marker) | Right 6 lanes (excluding marker) |
|---|---|---|
| Blot | GAPDH for S4a | GAPDH for S4b |

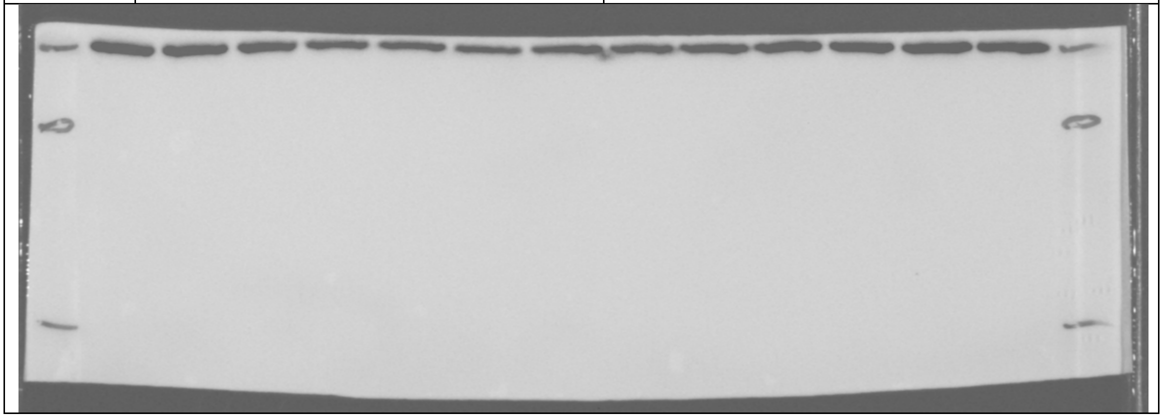

| | Left 6 lanes (excluding marker) | Right 6 lanes (excluding marker) |
|---|---|---|
| Blot | P53 for S4a | P53 for S4b |

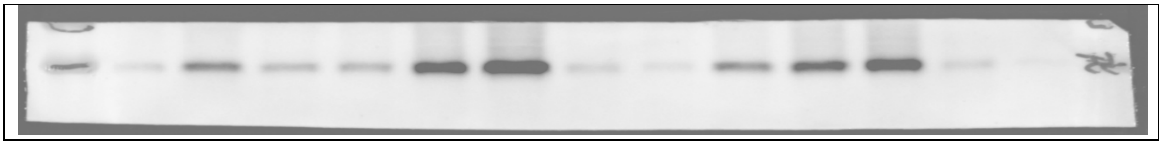

| | Left 6 lanes (excluding marker) | Right 6 lanes (excluding marker) |
|---|---|---|
| Blot | CHOP for S4a | CHOP for S4b |

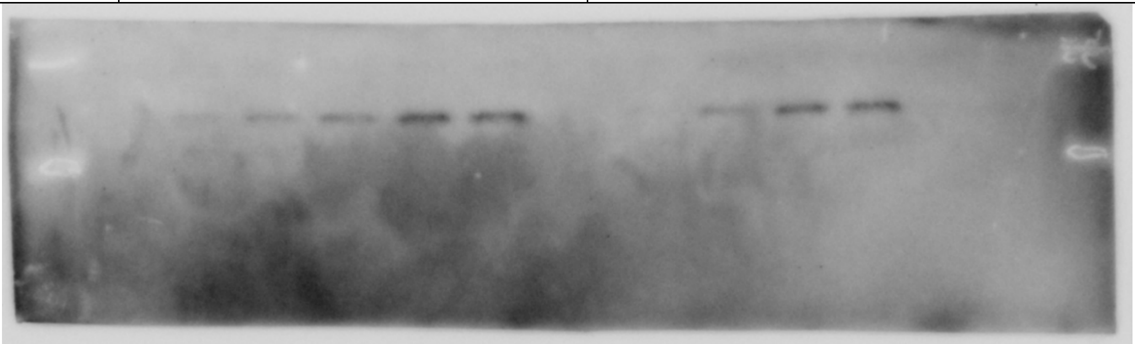

| | Left 6 lanes (excluding marker) | Right 6 lanes (excluding marker) |
|---|---|---|
| Blot | GAPDH for S4a | GAPDH for S4b |

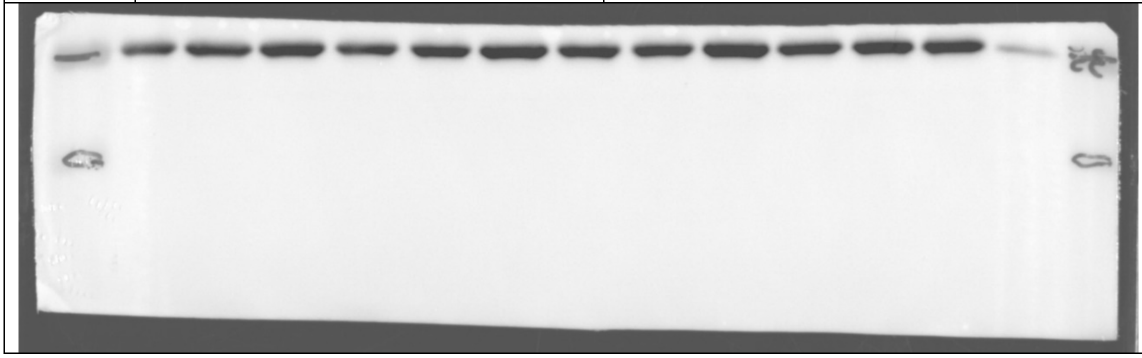

Uncropped blot for Figure S4c:

| | Lanes 8-12 (excluding marker) |
|---|---|
| Blot | IRE-1α |

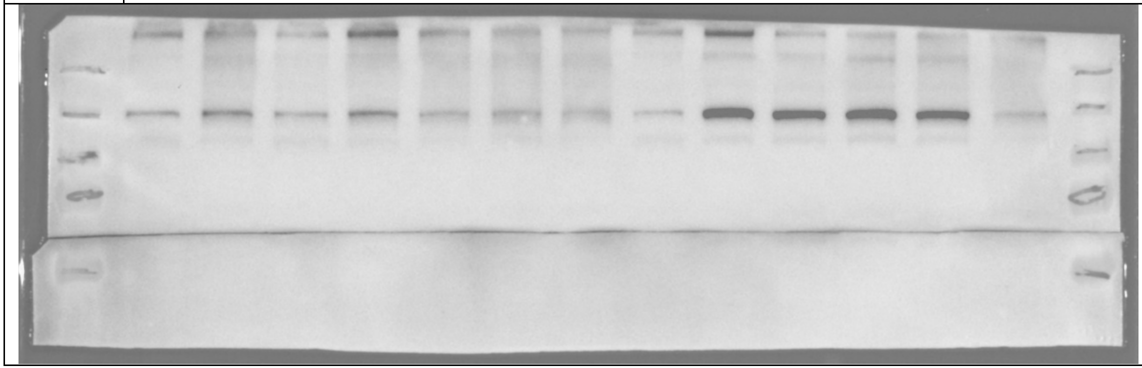

| | Lanes 8-12 (excluding marker) |
|---|---|

| Blot | p-IRE-1α |
| --- | --- |

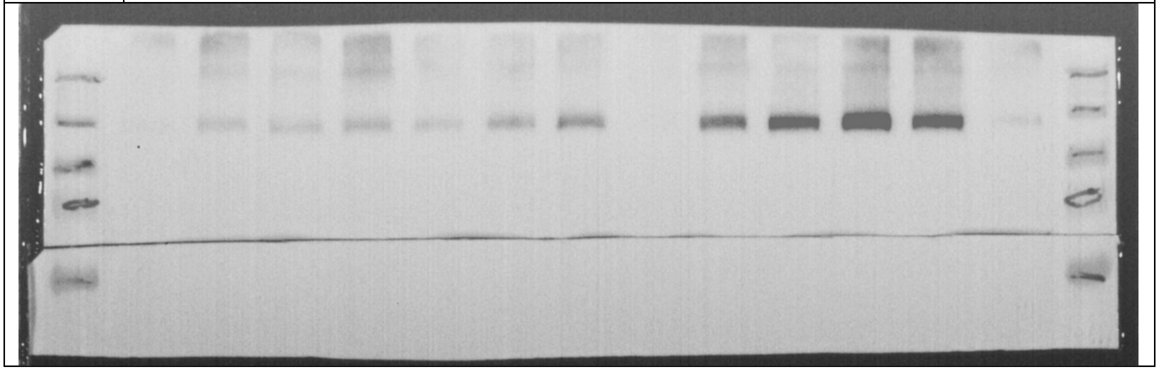

| | Lanes 8-12 (excluding marker) |
| --- | --- |
| Blot | GAPDH |

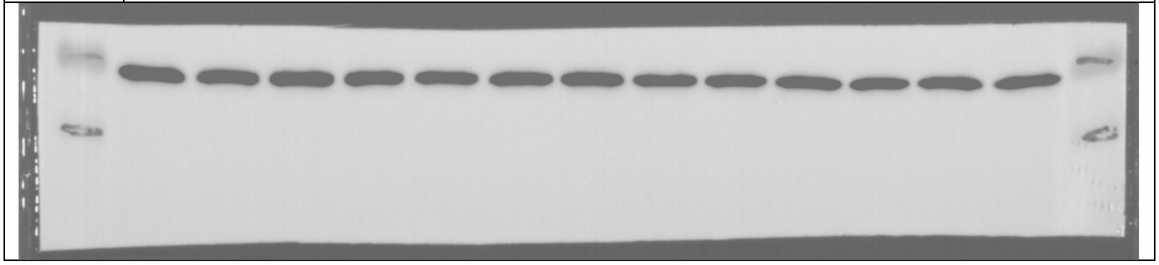
